# Selective transduction and photoinhibition of pre-Botzinger neurons that project to the facial nucleus in rats affect the nasofacial activity

**DOI:** 10.1101/2022.12.28.522095

**Authors:** MR Melo, A Wykes, AA Connelly, JK Bassi, SD Cheung, S McDougall, C Menuet, RAD Bathgate, AM Allen

**Affiliations:** Department of Anatomy & Physiology, University of Melbourne, Victoria, Australia; Florey Institute of Neuroscience and Mental Health, Victoria, Australia; Institut de Neurobiologie de la Méditerrané, INMED UMR1249, INSERM, Aix-Marseille Université, Marseille, France; Department of Biochemistry and Molecular Biology, University of Melbourne, Victoria, Australia.; Biological Optical Microscopy Platform (BOMP) - University of Melbourne, Victoria, Australia.

## Abstract

The preBötzinger Complex (preBötC), a key primary generator of the inspiratory breathing rhythm, contains neurons that project directly to facial nucleus (7n) motoneurons to coordinate orofacial and nasofacial activity. To further understand the identity of 7n-projecting preBötC neurons, we used a combination of optogenetic viral transgenic approaches to demonstrate that selective photoinhibition of these neurons affects mystacial pad activity, with minimal effects on breathing. These effects are altered by the type of anesthetic employed and also between anesthetised and conscious states. The population of 7n-projecting preBötC neurons we transduced consisted of both excitatory and inhibitory neurons that also send collaterals to multiple brainstem nuclei involved with the regulation of autonomic activity. We show that modulation of subgroups of preBötC neurons, based on their axonal projections, is a useful strategy to improve our understanding of the mechanisms that coordinate and integrate breathing with different motor and physiological behaviours. This is of fundamental importance, given that abnormal respiratory modulation of autonomic activity and orofacial behaviours have been associated with the development and progression of diseases.

## INTRODUCTION

Breathing is a complex behaviour requiring the coordination of multiple motor patterns in the face, upper airways, thorax and abdomen to enable gas exchange between the external and internal environment and maintenance of blood gas homeostasis (del Negro et al., 2018). Strict coordination of breathing with the activity of facial and upper airway muscles underlies the rhythmic orofacial behaviours, such as sniffing, sucking, licking, whisking, vocalization and swallowing, which are fundamental for ingestive and exploratory behaviour, and social interaction (Deschênes et al., 2016, 2015; Huff et al., 2022; Moore et al., 2014; Takatoh et al., 2022). Breathing-related information permeates the brain with respiratory modulation of suprapontine brain structures affecting cognitive function and emotional expression (Ashhad et al., 2022; Lavretsky and Feldman, 2021; Yackle et al., 2017). Similarly, coordination of this motor activity with autonomic nervous system activity produces phasic fluctuations of blood pressure (BP) and heart rate (HR) that are essential for optimizing cardiac efficiency and organ blood flow (Fisher et al., 2022; Menuet et al., 2020; O’Callaghan et al., 2020; Shanks et al., 2022).

The network of neurons coordinating breathing is distributed throughout the brainstem, with a major group responsible for generating inspiration located in the preBötzinger complex (preBötC) within the ventrolateral medulla oblongata (Smith et al., 1991). In rodents, the preBötC consists of a heterogeneous population of ∼3000 neurons with distinct, but intermingled, excitatory and inhibitory sub-groups (Ashhad and Feldman, 2020; del Negro et al., 2018). Anatomical studies showed that preBötC sends parallel excitatory and inhibitory projections to target nuclei throughout the brain, including to regions responsible for respiratory motor activity (Yang and Feldman, 2018).

As indicated by their extensive projections, preBötC neurons modulate many circuits in addition to those required for breathing activity. For example, preBötC neurons project monosynaptically to key medullary centers for autonomic regulation, and modulate cardio-respiratory coupling (Dempsey et al., 2017; Menuet et al., 2020, 2017). Photoinhibition of preBötC neurons produces the expected apnea, but also causes a significant reduction of sympathetic vasomotor activity, reduces respiratory modulation of BP and increases cardiac parasympathetic nerve activity to decrease HR and respiratory-sinus arrhythmia (RSA; Menuet et al., 2020). Neurons in the preBötC also modulate orofacial behaviours. Activation of the developing brain homeobox1 (Dbx1)-expressing preBötC neurons alters the networks responsible for the coordination of upper airways and swallowing, to decrease the probability of swallowing during inspiration (Huff et al., 2022). The preBötC also contributes to motor patterns that are necessary for exploratory behaviours, such as sniffing and whisking. In this case, a group of inhibitory preBötC neurons provide monosynaptic inputs onto the vibrissal premotor neurons in the intermediate reticular nucleus (vIRt) to facilitate synchronous whisking. In addition, preBötC contains facial premotor neurons that modulate nasal dilation and mystacial pad muscle activity to couple whisking to the breathing rhythm (Deschênes et al., 2016; Takatoh et al., 2022).

Understanding and identifying the preBötC subgroups that coordinate and integrate breathing with different motor and physiological behaviours is of fundamental importance, given that abnormal respiratory modulation of autonomic activity and orofacial behaviours have been associated with the development and progression of diseases (El-Omar et al., 2001; Huff et al., 2022; Menuet et al., 2017; Simms et al., 2009). Although several studies have attempted to understand the physiological function/s of each preBötC neuronal subpopulation, disturbances in breathing, that occur when preBötC neurons are either inhibited or activated, impact our understanding of whether the observed changes are due to the direct effects of preBötC neurons or alterations resulting from changes in the respiratory rhythm and blood gases (Cui et al., 2016; Huff et al., 2022; Menuet et al., 2020; Tan et al., 2008; Yackle et al., 2017). Furthermore, whilst substantial gains have been made in understanding the expression of selective molecular markers in sub-groups of preBötC neurons, it is rarely possible to reliably assign particular neurochemical signatures to unique functions (Ashhad and Feldman, 2020; del Negro et al., 2018; Yackle et al., 2017). For example, whilst Dbx1^+^SST^+^ predominantly affect the breathing pattern (Cui et al., 2016; del Negro et al., 2018; Ashhad and Feldman, 2020), some Dbx1^+^SST^+^ which also express the neurokinin-1 receptor (NK1R) and, in some cases, cadherin-9 (Cdh9+), promote generalized behavioural arousal (Yackle et al., 2017).

To further understand the organization of the preBötC, in this study, we have tested the hypothesis that the preBötC region is composed of segregated subgroups of output neurons, potentially driven by a separate group of rhythmogenic neurons, that modulate specific behaviours, such as orofacial muscle activity. If correct, we predict that selective photoinhibition of preBötC neurons projecting to the facial nucleus (7n) should modulate orofacial behaviour with minimal impact on respiratory motor activity. To test this hypothesis, we utilized a combinatorial viral transgenic approach with one virus providing cre-recombinase (Cre)-dependent expression of an optically-activated chloride channel, GtACR2, and another retrograde axonal delivery of Cre. We showed that selective transduction and photoinhibition of preBötC facial premotor neurons affects the mystacial pad activity whilst minimally affecting breathing.

## RESULTS

### Identification of different subpopulations of preBötC neurons based on their anatomical projection

To evaluate whether distinct subgroups of preBötC neurons could be distinguished based on their axonal projections, retrograde adeno-associated viruses (AAVrg), expressing either green fluorescent protein (GFP), or the red fluorophore, mCherry, were injected into 7n and the rostral ventrolateral medulla (RVLM), respectively (**Figure 1A**). Three weeks later, fluorophore expression was observed in preBötC neurons, with overlapping distribution (**Figure 1B**), albeit with the RVLM projecting cells tending to be more medial. Most cells only contained a single fluorophore, but a small number of double-labelled neurons were also identified (**Figure 1C**). Comparable results were obtained following the injection of Alexa Fluor 647-conjugated cholera toxin subunit B (CTB) and Alexa Fluor 555-conjugated CTB into 7n and RVLM (Figure 1-figure supplement **1A-D**). These results provide confidence that subgroups of preBötC neurons can be transduced based on their anatomical connection.

**Figure 1:**
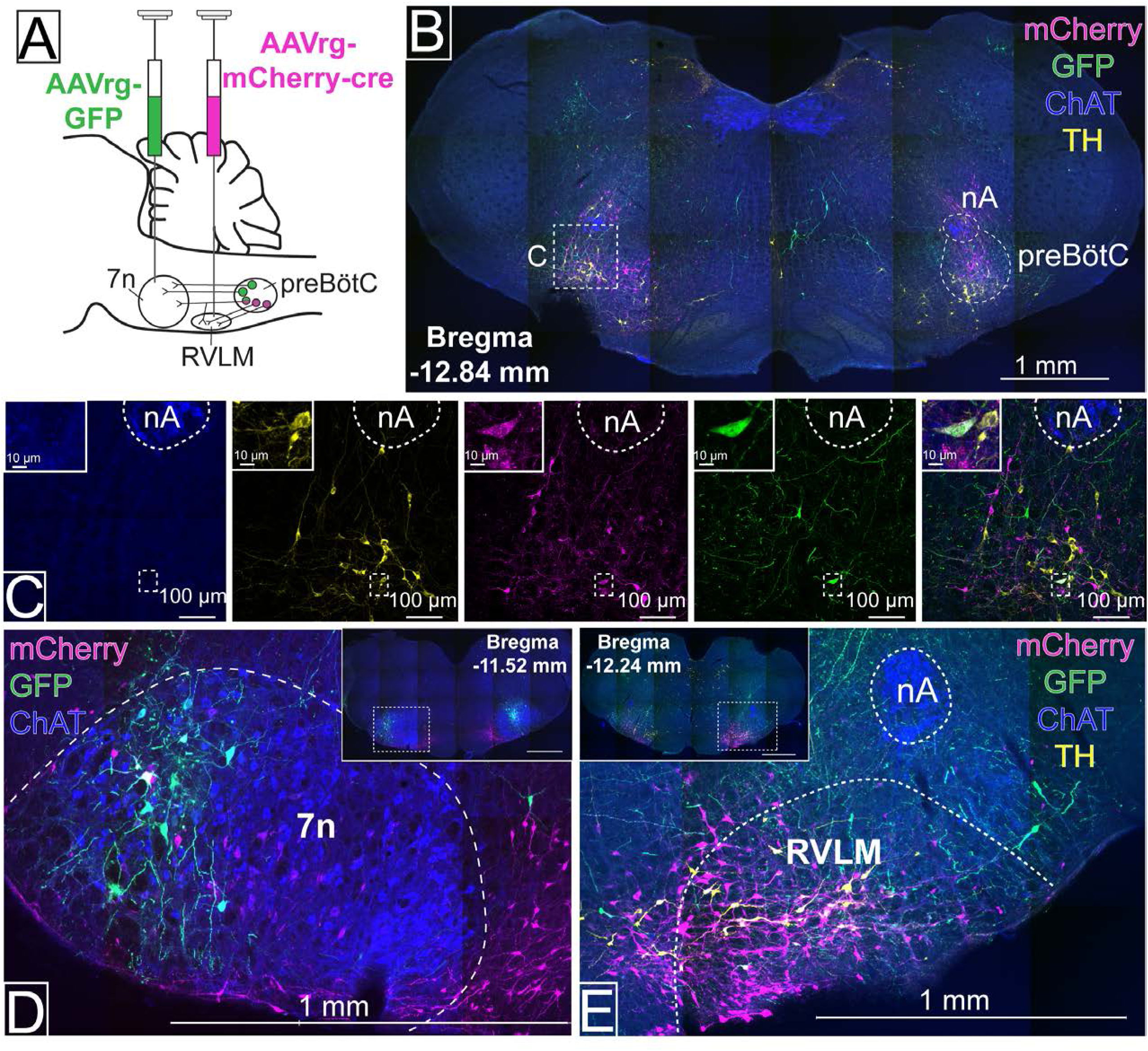
Distinct preBötC subpopulations project to the facial nucleus (7n) and rostral ventrolateral medulla (RVLM). (A) Schematic diagram showing the dual retrograde viral strategy. (B) Coronal section showing preBötC neurons projecting to 7n (green) and RVLM (red). Tyrosine hydroxylase (TH; yellow) and choline acetyltransferase (ChAT; blue) neurons are shown. (C) Transduced preBötC neurons, highlighted in the hashed box in (B), shown in higher magnification. The inset shows a preBötC neuron that projects to both 7n and RVLM. Local transduction shows the 7n (D) and RVLM (E) injection sites. Abbreviations: nA: nucleus ambiguus; The retrograde labelling of preBötC neurons from RVLM and 7n was also confirmed by injection of Alexa Fluor 647-conjugated cholera toxin subunit B (CTB) and Alexa Fluor 555-conjugated CTB into 7n and RVLM, as shown on Figure 1-Figure supplement 1. ***Figure supplement 1.*** Distinct preBötC neurons project to facial nucleus (7n) and rostral ventrolateral medulla (RVLM).

### Inhibition of preBötC neurons affects multiple physiological functions in urethane-anesthetized rats

To modulate the activity of selectively transduced neurons, we generated an adeno-associated virus (AAV) which enabled Cre-dependent (DIO) expression, under the control of the ubiquitous chicken β-actin/cytomegalovirus (CAG) promoter, of the light-activated chloride channel, GtACR2, fused to a portion of Kv2.1 to promote predominant somatic targeting (Lim et al., 2000), and an enhanced GFP (MuGFP; Scott et al., 2018); hereafter called AAV-DIO-GtACR2-MuGFP. This was co-injected with a Cre-expressing virus, AAVDJ8-Cre-TdTomato, in male Sprague Dawley rats (**Figure 2A**), to transduce all preBötC neurons in the region ventral and caudal to the tip of compact nucleus ambiguus. Immunohistochemical analysis showed that Cre recombination was effective and resulted in GtACR2-MuGFP expression in the preBötC (**Figure 2B**; Figure 2 – figure supplement **1A**). The Kv2.1 motif improved the neuronal expression of GtACR2, compared to GtACR2 alone (Menuet et al., 2020), with a clear definition of neuronal boundaries (**Figure 2C**). However, as previously demonstrated by other studies, the incorporation of the Kv2.1 motif into our viral construction did not prevent axonal expression of GtACR2-MuGFP (Mahn et al., 2018; Messier et al., 2018).At the caudal extent, a small proportion of GtACR2-MuGFP neurons also expressed parvalbumin, indicating likely transduction of some bulbospinal neurons of the rostral ventral respiratory group (rVRG; Figure 2 – figure supplement **1B**) and some vibrissa intermediate reticular nucleus (vIRt) neurons, that are located medially to nucleus ambiguus (Figure 2 – figure supplement **1C-F**). Control injections of AAV-DIO-GtACR2-MuGFP alone into preBötC did not produce any transduction (Figure 2 – figure supplement **2A-C**). Photoinhibition of these non-selectively transduced preBötC neurons in urethane-anesthetized rats produced very similar results to those described previously using a non-Cre-dependent approach (Menuet et al., 2020). This included immediate interruption of breathing and long-lasting apnea. Apnea did not last for the entire photoinhibition period, with inspiration resuming 9.3 s [95% CI: 6.3 to 12.3 s], after the beginning of the stimulus, albeit with a slower frequency and shorter dEMG amplitude during continued photoinhibition (**Figure 2D-F and M**). We also observed a biphasic BP response, with a depressor response during the apnea and a pressor response upon resumption of breathing (**Figure 2D, G and H**). We also observed a reduction in HR that lasted the entire period of photoinhibition, although it was not statistically different (**Figure 2I, J).**

**Figure 2:**
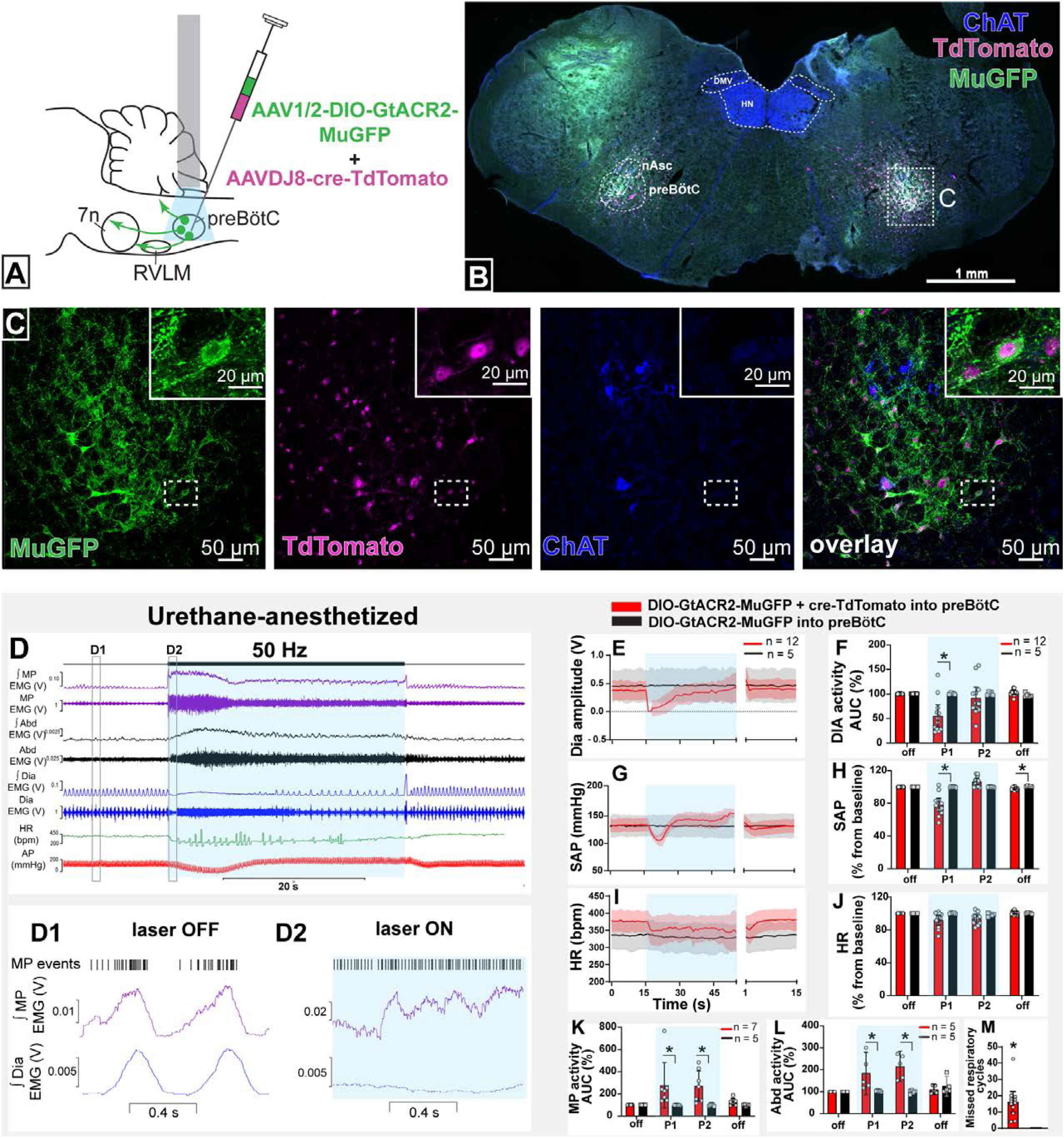
Effect on non-selective photoinhibition of preBötC on cardiovascular, respiratory and mystacial pad activity in urethane-anesthetized rats. (A) Schematic diagram showing the preBötC injection strategy. (B) Coronal section showing the expression of GtACR2-MuGFP and TdTomato in the preBötC, and choline acetyltransferase immunoreactivity (ChAT). Detailed maps showing the distribution of GtACR2-MuGFP expression are in Figure 2 *– figure supplement 1.* (C) Higher magnification confocal images of MuGFP and TdTomato in preBötC with the inset highlighting a double-labelled neuron. (D) Representative traces showing recordings of integrated (∫) and raw mystacial pad (MP) EMG, ∫ and raw abdominal muscle (Abd) EMG, ∫ and raw diaphragm (Dia) EMG, heart rate (HR), and arterial pressure (AP). Bilateral photoinhibition of preBötC, which occurs in the time highlighted by a blue box, produced increased, tonic MP and Abd activity, apnea, a biphasic change in AP, and decreased HR. Higher temporal resolution recordings of the periods highlighted by the hashed boxes are shown in (D1) and (D2), which show MP bursts as events, and triggered averages of ∫ MP and ∫ Dia EMG. Note that the inspiratory-related MP activity was interrupted by photoinhibition and MP EMG became increased and tonic. Group data showing mean (solid line) and 95% confidence intervals for (E) Dia amplitude, (G) systolic arterial pressure (SAP), (I) HR (bpm) before, during and after photoinhibition in GtACR2 expressing (red) and control (black) rats. Histograms showing group data for the effect of photoinhibition of preBötC in respiratory and cardiovascular parameters (F) Dia amplitude, (H) SAP, (J) HR, (K) MP activity, (L) Abd activity and (M) number of missed respiratory cycles; P1 and P2 refer to the initial period of photoinhibition, where apnea was complete, and the later period where breathing re-started respectively. The group data are presented as mean ± 95% CI; unpaired t-test or nonparametric Mann-Whitney test with multiple comparisons using the Bonferroni-Dunn method, *p<0.05. Abbreviations: nAsc: subcompact formation of the nucleus ambiguus; DMV: dorsal motor nucleus of the vagus; HN: hypoglossal nucleus. Control injections of AAV-DIO-GtACR2-MuGFP alone into preBötC are shown in Figure 2 *– figure supplement*. ***Figure supplement 1.*** Expression of GtACR2-MuGFP in rats co-injected with AAV-DIO-GtACR2-MuGFP and AAVDJ8-Cre-TdTomato in the preBötC. ***Figure supplement 2.*** Control experiments with the injection of AAV-DIO-GtACR2-MuGFP alone into preBötC. **Source data 1.** Source data and statistics for Figure 2

Orofacial muscle activity, measured from electrodes inserted into the mystacial pad near the rostral tip of the snout - the site of origin of the muscle *nasolabialis profundus* (Haidarliu et al., 2010), showed activity during the pre-inspiratory phase of the respiratory cycle, associated with the initial phase of vibrissae protraction (Deschênes et al., 2016). With non-selective photoinhibition of preBötC neurons, the mystacial pad (MP) activity increased in amplitude and shifted and became tonic (**Figure 2D, D1, D2** and **K**) with loss of inspiratory-related modulation. The onset of these effects correlated with apnea. When breathing resumed, during continued photoinhibition, the inspiratory-related MP activity also recovered, with increased amplitude, for the remainder of the stimulation period. Photoinhibition of the preBötC also immediately increased and produced tonic activity of the abdominal (Abd) muscle that lasted for the apnea duration (**Figure 2D, L)**. In some rats, rhythmic expiratory activity was observed with the return of dEMG inspiratory activity. Our results show that preBötC exerts a complex influence on orofacial activity as well as the timing of active expiration, potentially via the parafacial respiratory group (pFRG).

No changes in any of these parameters were observed in control rats injected with only AAV-DIO-GtACR2-MuGFP (**Figure 2E-M** and Figure 2 – figure supplement **2D**).

### Photoinhibition of preBötC neurons that project to the facial nucleus

We injected AAVrg-mCherry-Cre into the lateral and dorsal lateral border of the facial nucleus. This region is known to contain respiratory-modulated facial motoneurons that receive projections from the preBötC (Deschênes et al., 2016; Takatoh et al., 2013), with the lateral most edge of the facial nucleus containing motoneurons that regulate pre-inspiratory phase activity of the facial *nasolabialis profundus* muscle (NLP) to cause vibrissae protraction (Deschênes et al., 2016). In the same surgery, we also injected AAV-DIO-GtACR2-MuGFP into the preBötC with the aim of selectively transducing preBötC neurons projecting to the facial nucleus (preBötC→7n; **Figure 3A**).

**Figure 3:**
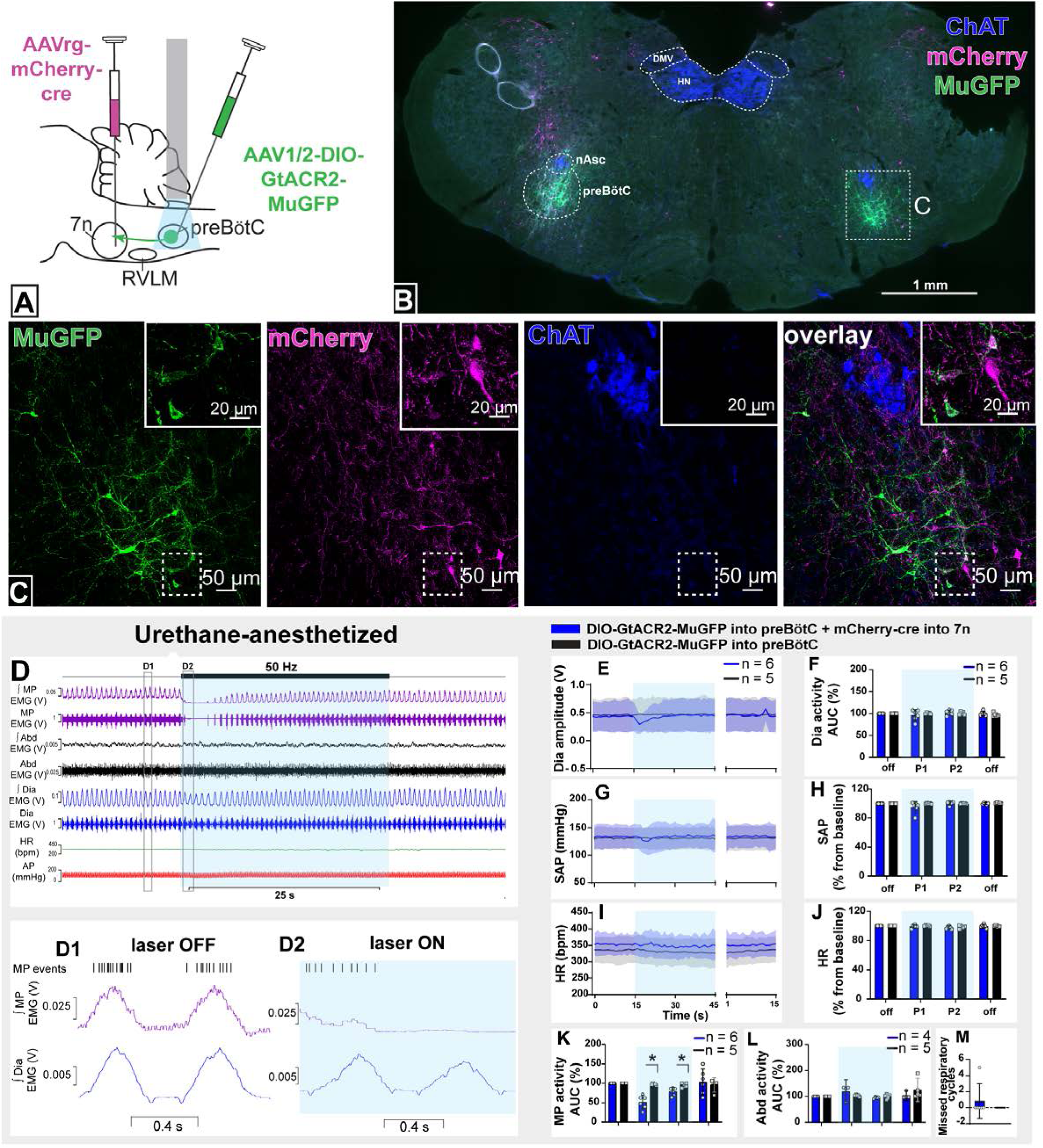
Effect of selective inhibition of preBötC→7n neurons on cardiovascular, respiratory and nasofacial activity in urethane-anesthetized rats. (A) Schematic diagram showing the injection protocol for selective transduction of preBötC→7n neurons. (B) Coronal section showing the expression of GtACR2-MuGFP and mCherry into the preBötC and immunohistochemistry for choline acetyltransferase (ChAT). Detailed maps showing the distribution of the expression of GtACR2-MuGFP are shown in Figure 3 *– figure supplement 1*. (C) Higher magnification confocal image showing the co-localization of MuGFP with mCherry in neurons of preBötC. (D) Representative trace showing the integrated (∫) and raw mystacial pad (MP) EMG, ∫ and raw abdominal muscle (Abd) EMG, ∫ and raw diaphragm (Dia) EMG, heart rate (HR), and arterial pressure (AP). Bilateral selective photoinhibition of preBötC→7n neurons, highlighted with the blue box, decreased the overall activity, and interrupted the inspiratory-related MP activity, with minimal effects on respiratory, cardiovascular or abdominal muscle activity. Higher temporal resolution recordings of the periods highlighted by the hashed boxes are shown in (D1) and (D2), which show MP bursts as events, and triggered averages of ∫ MP and ∫ Dia EMG. Note that the overall MP activity decreased, even in the absence of the interruption of inspiratory activity, the inspiratory-related MP activity ceased; Group data showing mean (solid line) and 95% confidence intervals for (E) Dia amplitude, (G) systolic arterial pressure (SAP) and (I) HR (bpm) before, during and after photoinhibition in selective GtACR2 expressing (blue) and control (black) rats. Histograms showing group data for the effect of photoinhibition of preBötC on respiratory and cardiovascular parameters (F) Dia amplitude, (H) SAP, (J) HR, (K) MP activity, (L) Abd activity and (M) number of missed respiratory cycles; P1 and P2 refer to the initial period of photoinhibition, where there is a small decrease in Dia amplitude and the later period respectively. Group data are presented as mean ± 95% CI; unpaired t-test or nonparametric Mann-Whitney test with multiple comparisons using the Bonferroni-Dunn method, *p<0.05. The period of photoinhibition of preBötC neurons is depicted by blue shading. Abbreviations: DMV: dorsal motor nucleus of the vagus; nAsc: subcompact formation of the nucleus ambiguus; HN: hypoglossal nucleus. ***Figure supplement 1.*** Expression of GtACR2-MuGFP in selective preBötC→7n transduced rats. **Source data 1.** Source data and statistics for Figure 3

This combinatorial approach resulted in GtACR2-MuGFP expression in a restricted, defined subgroup of preBötC neurons mostly located ventral to the extension of the subcompact nucleus ambiguus (**Figure 3B,C** and Figure 3 – figure supplement **1A**). Some of the transduced neurons were also found intermingled with parvalbumin-expressing rVRG neurons, but no co-localization between MuGFP and parvalbumin was observed (Figure 3 – figure supplement **1B**). A very small number of non-parvalbumin neurons, as well as sparse axonal labeling, was also observed medial to the nucleus ambiguus, in the region of the vIRt (Figure 3 – figure supplement **1C-F**).

In urethane-anesthetized rats, photoinhibition of preBötC→7n neurons significantly reduced the amplitude of the mystacial pad activity for 7.3 s [95% CI: 0.5 to 14.2 s], with complete abolition of the inspiratory-related activity of the mystacial pad in half of the rats (Figures **3D, D1, D2, K**). In some rats, we observed a small decrease in dEMG amplitude and respiratory frequency, but this was not statistically significant for the cohort (**Figure 3E, F and G**). A slight reduction in BP was also observed at the beginning of the photoinhibition, but again it was not statistically significant (Figures **3G,H**). Photoinhibition of preBötC→7n did not affect HR or Abd activity (Figures **3I,J and L) -**. These results clearly identify a subgroup of preBötC neurons providing inspiratory modulation of facial motoneurons that innervate extrinsic protractor muscles of the mystacial pad, which appear to be largely independent of those that drive inspiratory activity to the diaphragm, abdominal muscles and autonomic nervous system.

### Anatomical and neurochemical characterization of transduced preBötC neurons

To characterize the anatomical distribution and neurochemical phenotype of preBötC→7n neurons, we combined immunohistochemistry and RNAscope, to identify mRNA expression. Following non-selective transduction of preBötC, GtACR2-MuGFP expression occurred between 12.5 and 13.92 mm caudal to Bregma (**Figure 4A,B).** We examined expression in detail in three rats and counted 1027 transduced neurons [95% CI: 769 to 1286 neurons] in each rat. Of these, 60.8% [95% CI: 59.2 to 62.4 %] neurons expressed the vesicular GABA transporter (VGAT), and 32.3% [95% CI: 30.5 to 34.1] the vesicular-glutamate transporter 2 (VGlut2; **Figure 4A-D**). The preBötC→7n transduced neurons were more restricted and located more caudally, between 13 and 13.76 mm caudal to Bregma (**Figure 4E,F**). In three rats, we counted 97 neurons transduced neurons [95% CI: 32 to 162 neurons] in each rat. As expected from previous reports (Yang and Feldman, 2018), a similar proportion of transduced preBötC→7n neurons expressed VGAT 38.6% [95% CI: 35.6 to 41.6 %] or VGlut2 38% [95% CI: -7 to 82.8%](**Figure 4E-H**).

**Figure 4:**
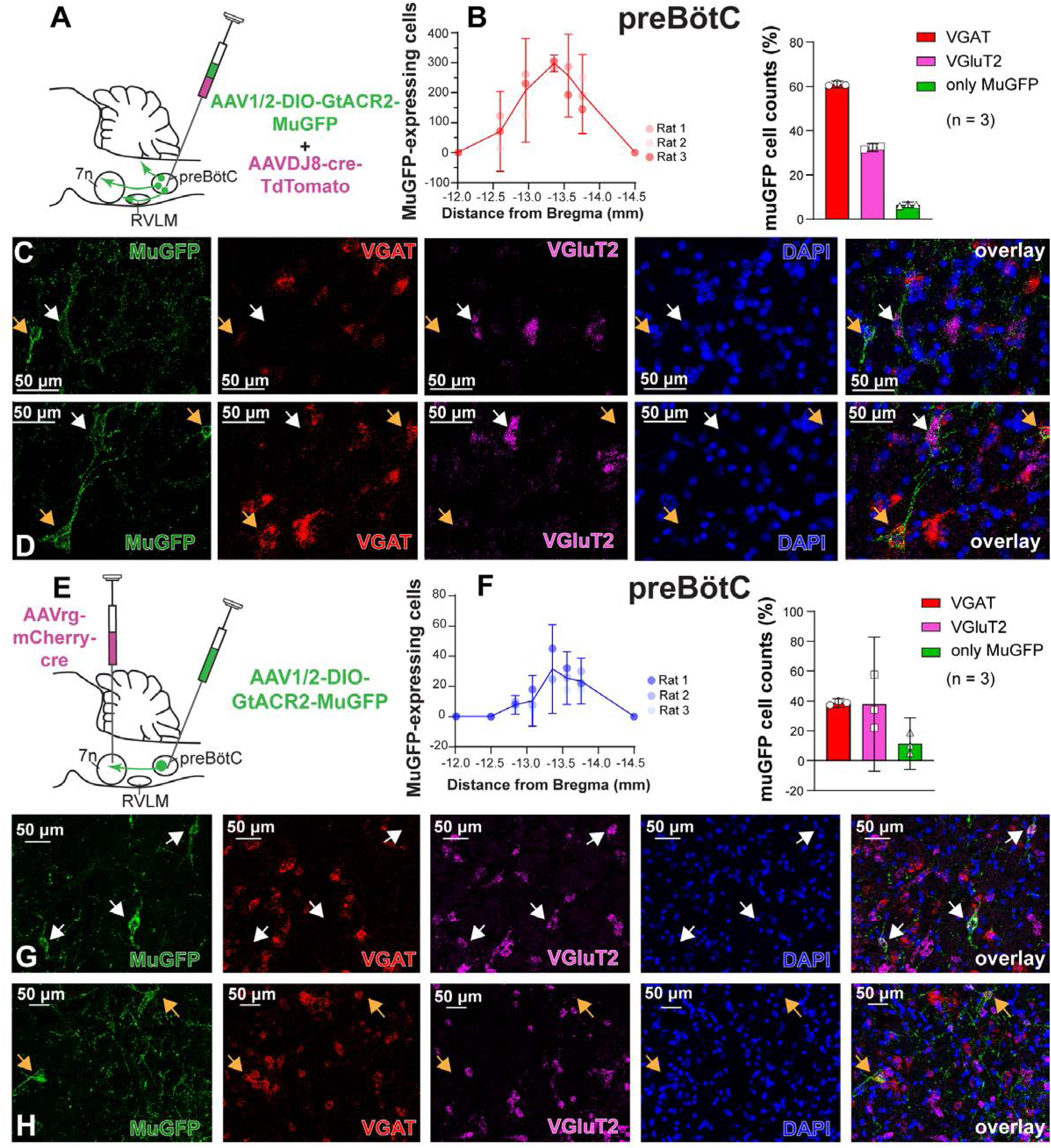
Excitatory and inhibitory preBötC neurons are transduced by either non-selective or selective transfection of preBötC→7n neurons. (A) Schematic diagram showing injection protocol for non-selective transduction of preBötC neurons. (B) Quantification of the total number of MuGFP-expressing neurons, plotted as the distance from Bregma (mm). The histograms show the number of transduced cells that co-expressed mRNA for *VGAT* or *VGluT2*. The results are presented as mean ± 95% CI. (C-D) *In situ hybridization* showing the co-expression of MuGFP (green), with mRNA for VGAT (red) or VGluT2 (magenta) in preBötC. Nuclei are labelled in blue (DAPI). The yellow arrows highlight *VGAT* neurons, and the white arrows highlight *VGluT2* neurons. (E) Schematic diagram showing the injection protocol for selective transduction of preBötC→7n neurons. (F) Quantification of the total number of MuGFP-expressing neurons, plotted as the distance from Bregma (mm). Note the slight trend towards a more caudal distribution. The histograms show the number of transduced cells that co-expressed mRNA for *VGAT* or *VGluT2*. The results are presented as mean ± 95% CI. (G-H) *In situ hybridization* showing the co-expression of MuGFP (green), with mRNA for VGAT (red) or VGluT2 (magenta) in preBötC→7n neurons. Nuclei are labelled in blue (DAPI). The yellow arrows highlight *VGAT* neurons, and the white arrows highlight *VGluT2* neurons.

### Distribution of the axonal projections of preBötC neurons projecting to the facial nucleus

Photoinhibition of preBötC→7n neurons produced a small effect on breathing and BP, although not statistically significant across the cohort. As suggested by our initial retrograde labeling experiments, where we observed a small number of preBötC with axon collaterals to both 7n and RVLM, we wondered whether collateralization might explain the small effects of photoinhibition of preBötC→7n neurons on non-facial motor outputs. Non-selective transduction of preBötC neurons resulted in a dense GtACR2-MuGFP-axonal expression in multiple brainstem nuclei (**Figure 5**), including the nucleus ambiguus (**Figure 5B**); RVLM (**Figure 5C**); Bötzinger Complex (BötC) (**Figure 5C**); hypoglossal nucleus (**Figure 5D**); nucleus of the solitary tract (NTS) (**Figure 5F**); 7n (**Figure 5E**); rVRG (Figure 2 – supplement **1B**) and vIRt (Figure 2 – supplement **2B-E**).

**Figure 5:**
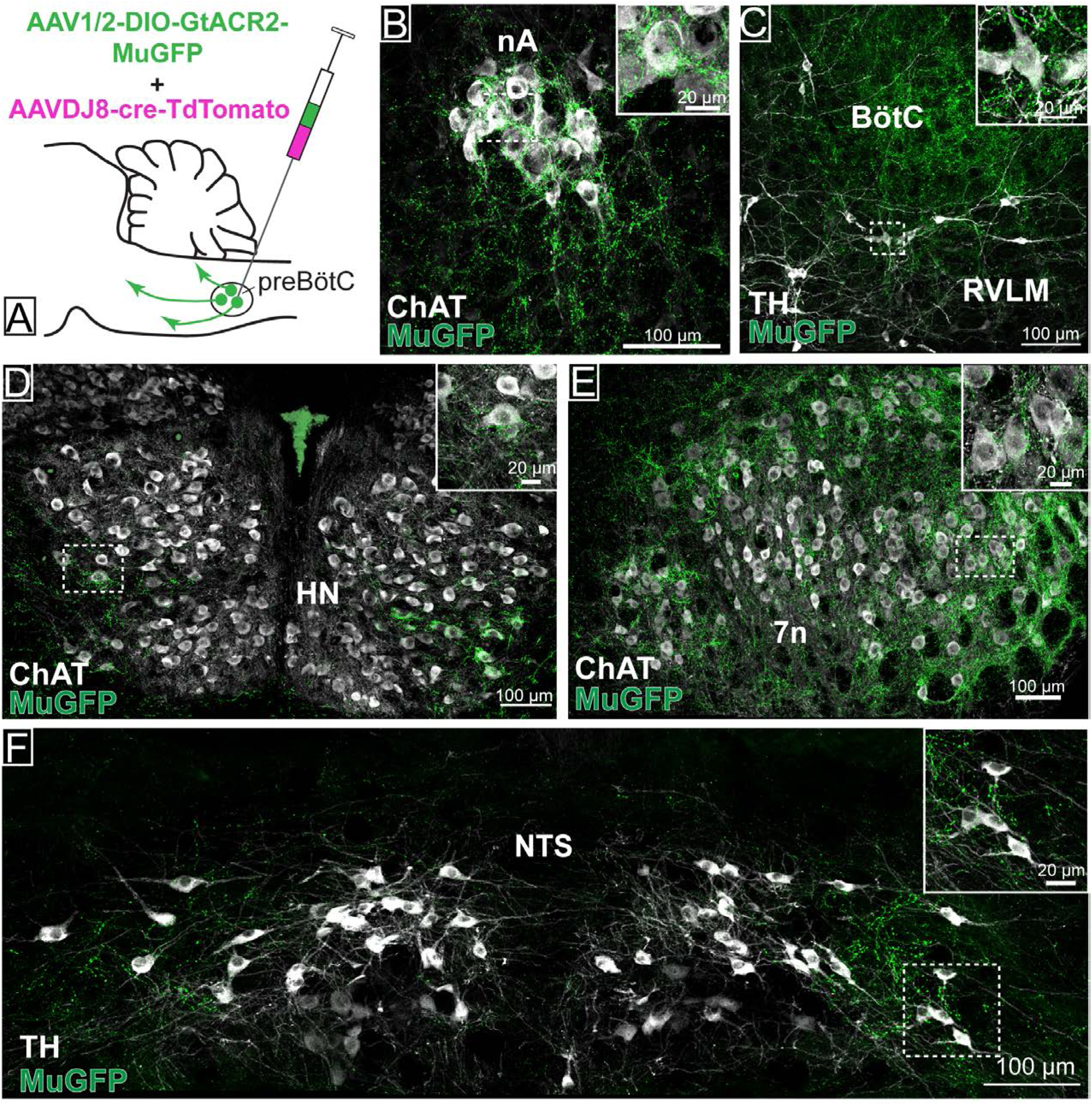
Distribution of GtACR2-MuGFP-expressing axons in multiple brainstem nuclei following the non-selective transduction of preBötC neurons. (A) Schematic diagram showing the protocol for non-selective transduction of preBötC neurons. Confocal microscopy images demonstrate MuGFP expression in axon in the (B) nucleus ambiguus. (C) rostral ventrolateral medulla and Bötzinger Complex, (D) hypoglossal nucleus, (E) facial nucleus and (F) nucleus of the solitary tract. Higher magnification images of the hashed-boxed regions are shown in the upper right corner of the lower magnification image. ChAT: choline acetyltransferase; TH: tyrosine hydroxylase.

Strong axonal labelling was observed in the 7n, particularly along the dorsal and lateral edge of the nucleus following transduction of preBötC→7n neurons (**Figure 6**). In these animals, we also observed sparse GtACR2-MuGFP-axonal expression in the nucleus ambiguus (**Figure 6B**); BötC (**Figure 6C**); RVLM (**Figure 6C**); hypoglossal nucleus (**Figure 6D**); NTS (**Figure 6F**); rVRG (Figure 2 – supplement **1D**) and vIRt (Figure 2 – supplement **2F-I**).

**Figure 6:**
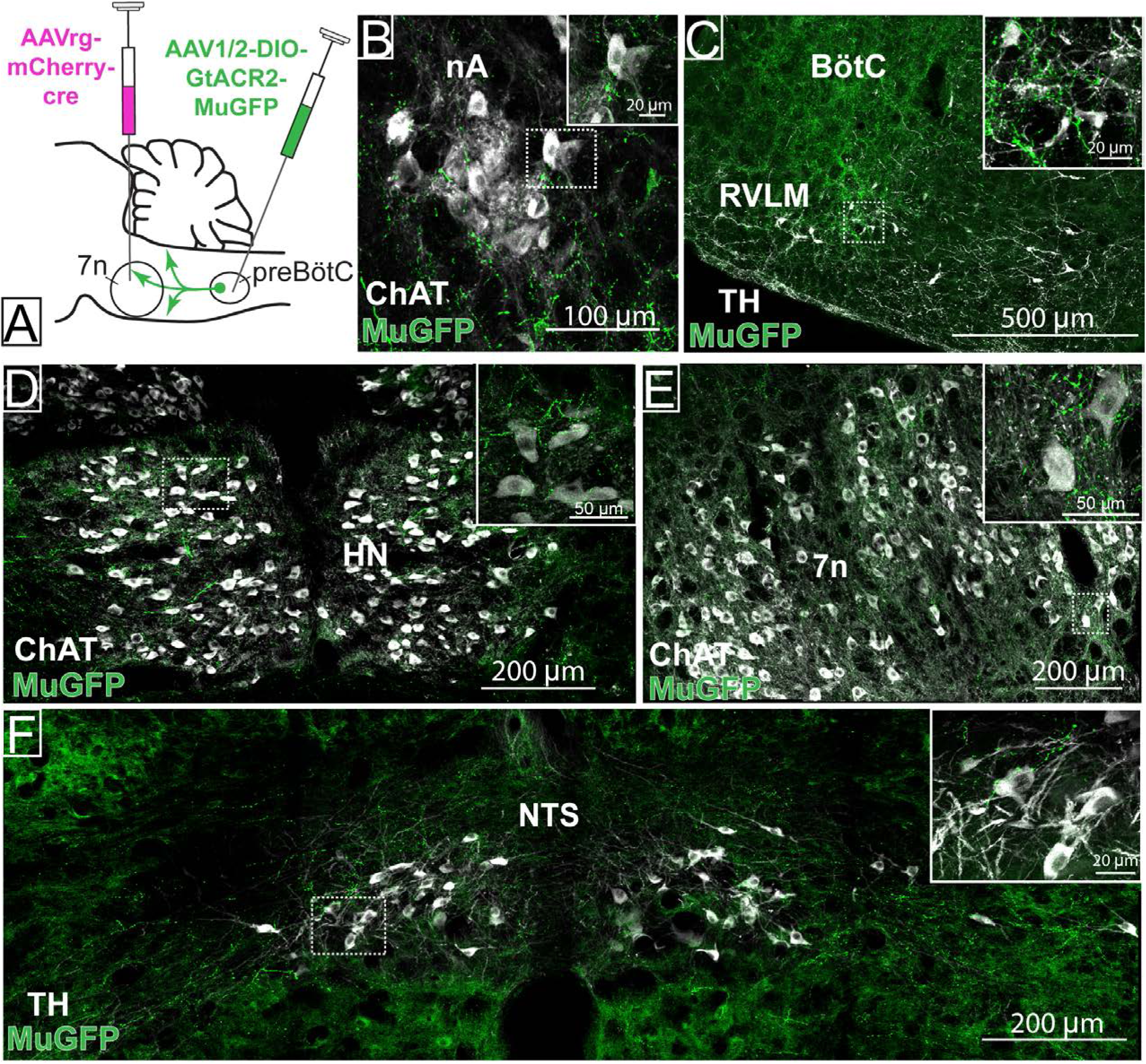
Distribution of selective preBötC→7n GtACR2-MuGFP-expressing axonal projections. (A) Schematic diagram showing the injection protocol for selective transduction of preBötC→7n neurons. Confocal microscopy images highlighting the expression of MuGFP in axons in the (B) nucleus ambiguus; (C) rostral ventrolateral medulla and Bötzinger Complex; (D) hypoglossal nucleus; (E) facial nucleus and (F) nucleus of the solitary tract. Higher magnification images of the hashed-boxed regions are shown in the upper right corner of the lower magnification image. ChAT: choline acetyltransferase; TH: tyrosine hydroxylase.

These results indicate widespread collateralization of preBötC neurons into distinct areas of the brainstem. Whether this is the result of single neurons projecting to many targets or many neurons projecting to a small number of targets remains to be investigated. We speculate that this anatomical organization may contribute to the complex synchronization between respiratory, cardiovascular, orofacial, and potentially other physiological, functions.

### Selective inhibition of preBötC neurons projecting to 7n in conscious rats

The rhythmic orofacial activities, such as whisking and sniffing, require a hierarchical organization of brainstem circuitry that involves the whisking premotor neurons in the vIRt and the preBötC respiratory-related premotor neurons (Deschênes et al., 2016; Takatoh et al., 2022). The activity of the whisking premotor neurons in vIRt is strongly affected by anesthesia and animal state (Deschênes et al., 2016, 2015). Here we evaluated whether the response to photoinhibition of preBötC neurons was also affected by the type of anesthesia and the animals’ state. Non-selective preBötC and selective preBötC→7n photoinhibition were performed under surgical ketamine/medetomidine anesthesia, during initial recovery to consciousness after reversal of anesthesia with atipamezole (1 mg/kg, i.p.), and 1-2 h after reversal of anesthesia.

### Surgical ketamine/medetomidine anesthesia

In contrast to the urethane-anesthetized rats, during surgical ketamine/medetomidine anesthesia the mystacial pad activity was minimal and non-rhythmic (Figure 7 – figure supplement **1B, B1,L, L1**). Photoinhibition of preBötC neurons induced long-lasting apnea without affecting mystacial pad activity or HR (Figure 7 – figure supplement 1 A-J). Photoinhibition of preBötC→7n neurons produced minimal effects on any parameter (Figure 7 – figure supplement **1K-T**). In one out of 5 rats, we observed a short apnea (3.9 s) (Figure 7 – figure supplement **1S**).

**Figure 7:**
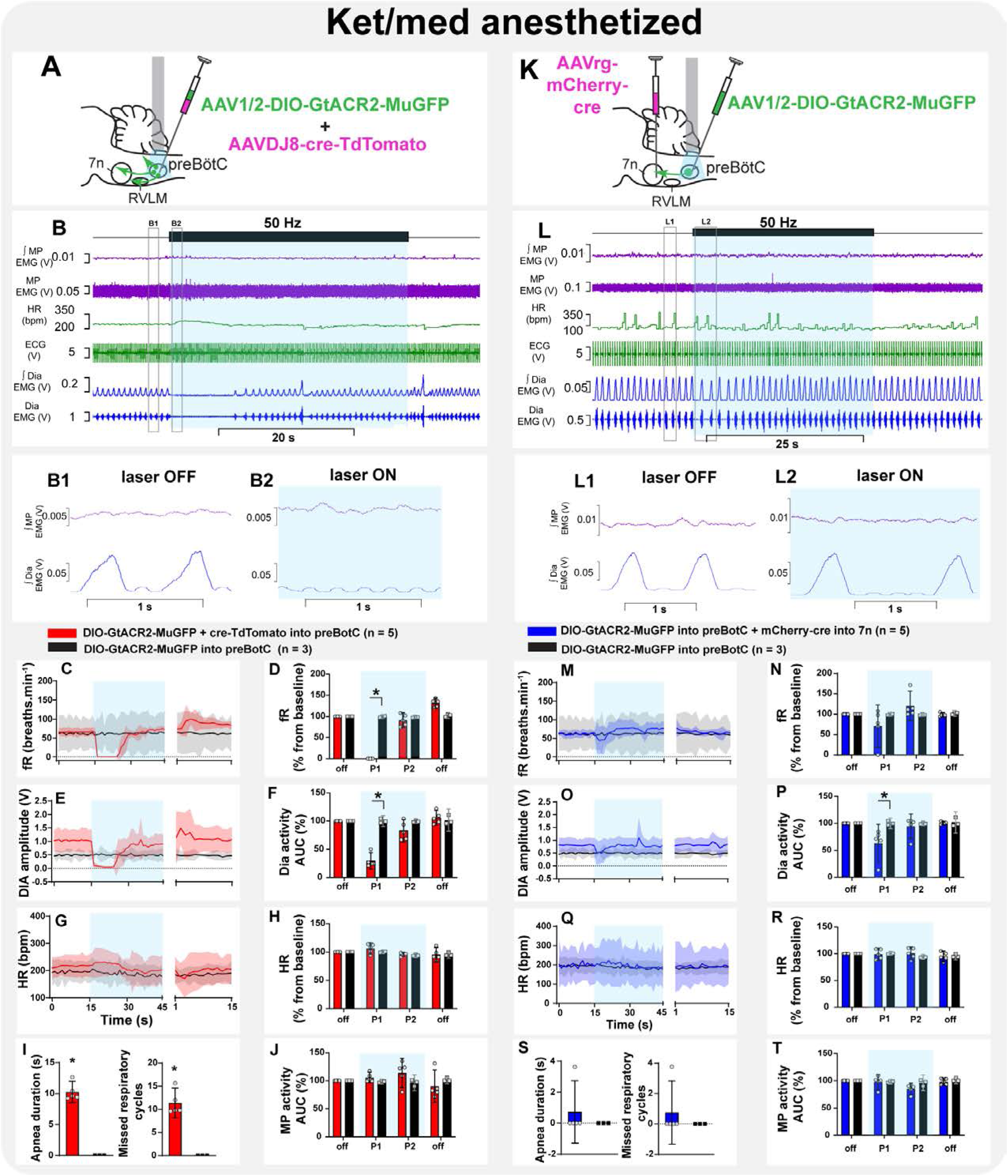
Effect of selective inhibition of preBötC→7n neurons on mystacial pad activity is state-dependent. Schematic diagrams showing the injection protocols for non-selective transduction of preBötC neurons (A) and selective transduction of preBötC→7n neurons (K). Representative trace showing the integrated (∫) and raw mystacial pad (MP) EMG, heart rate (HR), ECG, and ∫ and raw diaphragm (Dia) EMG in the initial phase of reversal of ketamine/medetomidine-anesthesia in rats with non-selective (B) and selective (L) preBötC neuron transduction. The period of photoinhibition is highlighted with the blue box. Higher resolution recordings of the hashed boxes showing inspiratory modulation of MP activity during baseline (B1, L1). Photoinhibition of non-selectively transduced preBötC neurons induced apnea and interrupted inspiratory-related MP activity, which increased and become tonic for the apnea period (B2). Photoinhibition of preBötC→7n neurons interrupted the inspiratory-related activity of MP, even in the absence of apnea (L2). Group data showing mean (solid line) and 95% confidence intervals for (C and M) respiratory frequency – fR, (E and O) Dia amplitude, (G and Q) and HR (bpm) before, during and after photoinhibition in non-selective (red), selective GtACR2 expressing (blue) and control (black) rats. Histograms showing group data for the effect of photoinhibition of preBötC on respiratory and cardiovascular parameters (D and N) fR, (F and P) Dia amplitude, (H and R) HR, (I and S) apnea duration and number of missed respiratory cycles, and (J and T) MP activity.; P1 and P2 refer to the initial period of photoinhibition Group data are presented as mean ± 95% CI; unpaired t-test or nonparametric Mann-Whitney test with multiple comparisons using the Bonferroni-Dunn method, *p<0.05. The photoinhibition of preBötC neurons is depicted by blue shading. The effects of non-selective and selective photoinhibition of preBötC in ketamine/medetomidine-anesthetized rats are shown in Figure 7 *– figure supplement 1*. ***Figure supplement 1:*** Effect on non-selective and selective inhibition of preBötC on mystacial pad activity under ketamine/medetomidine anesthesia. **Source data 1.** Source data and statistics for Figure 7

### Early recovery phase

During this phase, respiratory rate and HR gradually increased, and the inspiratory modulation of the mystacial pad activity resumed (**Figure 7B, B1,L, L1**). Spontaneous opening of the nares and whisker motion were visualized. Photoinhibition of preBötC immediately induced apnea, with mystacial pad activity becoming tonic, with no obvious inspiratory modulation (**Figure 7B, B2, C-F,** and **I**). The amplitude of the mystacial pad activity increased in all five rats, but this was not statistically significant (**Figure 7J**). With the resumption of breathing, mystacial pad activity returned to baseline, even when this occurred during continued light delivery. The HR was not affected (**Figure 7G,H**). Selective inhibition of preBötC→7n neurons did not induce apnea, although a slight reduction in respiratory frequency occurred (**Figure 7M-P** and **S).** In contrast to observations under the urethane anesthesia, photoinhibition of preBötC→7n neurons increased the overall mystacial pad activity during the first 5.1 s [95% CI: 2.9 to 7.4 s] from the onset of the stimulus and interrupted the inspiratory-related modulation (**Figure 7L, L1, L2** and **T)** and **S**). No changes were observed in HR (**Figure 7Q,R**).

### Conscious phase

Non-selective photoinhibition of preBötC neurons induced apnea and did not affect HR(Figure 8B, C-H). The amplitude of mystacial pad activity did not change, but became tonic (**Figure 8B, B1, B2** and **J**). In contrast, in the initial period of selective photoinhibition of preBötC→7n neurons, the inspiratory-modulated rhythm of mystacial pad activity was disrupted, and displayed a more tonic pattern (**Figure 8L, L1, L2** and **T)**. Diaphragm EMG was not affected, except for the induction of a very short apnea in two out the 5 rats, but respiratory frequency decreased throughout the entire stimulation period (**Figure 8L-P, and S**). Inhibition of preBötC→7n neurons did not affect HR (**Figure 8Q,R).**

**Figure 8:**
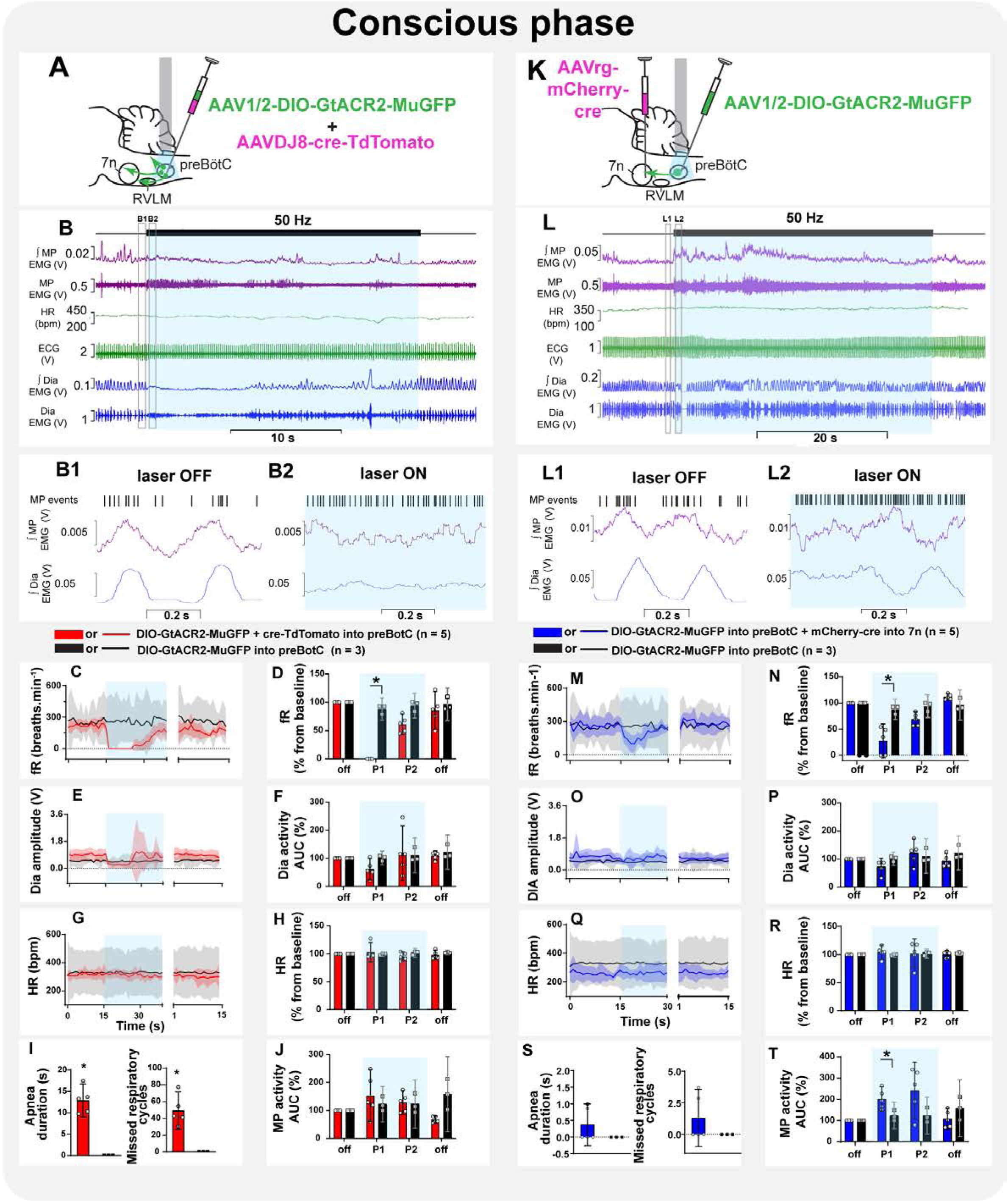
Inspiratory-related activity of mystacial pad is interrupted by the selective inhibition of preBötC to 7n neurons in conscious rats. Schematic diagrams showing the injection protocols for non-selective transduction of preBötC neurons (A) and selective transduction of preBötC→7n neurons (K). Representative trace showing the integrated (∫) and raw mystacial pad (MP) EMG, heart rate (HR), ECG, and ∫ and raw diaphragm (Dia) EMG after recovery from ketamine/medetomidine-anesthesia in rats with non-selective (B) and selective (L) preBötC neuron transduction. The period of photoinhibition is highlighted with the blue box. Higher resolution recordings of the hashed boxes showing inspiratory modulation of MP activity during baseline (B1, L1) and the initial phase of photoinhibition (B2, L2). The MP EMG is more variable in conscious rats, due to exploratory and other behaviors. Photoinhibition of non-selectively transduced preBötC neurons interrupted inspiratory-related MP activity, which increased and became tonic for the apnea period (B2). Photoinhibition of preBötC→7n neurons increased tonic activity of MP, which was mostly out of phase with respiration (L2).). Group data showing mean (solid line) and 95% confidence intervals for (C and M) respiratory frequency – fR, (E and O) Dia amplitude, (G and Q) and HR (bpm) before, during and after photoinhibition in non-selective (red), selective GtACR2 expressing (blue) and control (black) rats. Histograms showing group data for the effect of photoinhibition of preBötC on respiratory and cardiovascular parameters (D and N) fR, (F and P) Dia amplitude, (H and R) HR, (I and S) apnea duration and number of missed respiratory cycles, and (J and T) MP activity; P1 and P2 refer to the initial period of photoinhibition. Group data are presented as mean ± 95% CI; unpaired t-test or nonparametric Mann-Whitney test with multiple comparisons using the Bonferroni-Dunn method, *p<0.05. The photoinhibition of preBötC neurons is depicted by blue shading. **Source data 1.** Source data and statistics for Figure 8

## DISCUSSION

We employed a combinatorial Cre-dependent approach to transduce a sub-group of preBötC neurons based on their axonal projections to the facial nucleus. The preBötC is the kernel for breathing (Smith et al., 1991), but also acts as a master oscillator controlling cardiovascular and orofacial activities (del Negro et al., 2018; Deschênes et al., 2016; Huff et al., 2022; Takatoh et al., 2022), and our study aimed to determine whether these functions involved independent sub-groups of preBötC neurons. We validated our approach by co-injections of a Cre virus and the Cre-dependent anion channel virus – non-selective preBötC transduction. In agreement with previous results (Menuet et al., 2020) non-selective photoinhibition of preBötC neurons induced apnea, bradycardia and biphasic effects on BP. We also observed loss of inspiratory modulation of mystacial pad activity, with increased tonic activity, except in rats anesthetized with ketamine/medetomidine where there was no ongoing mystacial pad activity. By contrast, selective photoinhibition of preBötC→7n neurons had minimal effect on breathing or cardiovascular activity, but altered mystacial pad activity with the effect altered by the type of anesthetic or state. The preBötC→7n neurons showed a restricted anatomical distribution within the preBötC, but sent substantial collateral projections to several areas in the brainstem involved with the cardiorespiratory regulation. We conclude that these collaterals are functionally active and responsible for the small effects of photoinhibition of preBötC→7n neurons on BP, HR and respiratory frequency. Both selective preBötC→7n neuron transduction and non-selective preBötC transduction resulted in transgene expression in both GABAergic and glutamatergic neurons.

The observation that photoinhibition of preBötC→7n neurons significantly affects mystacial pad activity, without affecting breathing, BP or HR, clearly demonstrates that modulation of preBötC subgroups of neurons based on their axonal projections is a useful strategy.

It allows an understanding of the anatomical distribution and neurochemical phenotype of subgroups of preBötC neurons. It also enables assessment of the physiological relevance of these subgroups without interference and potentially the development of secondary physiological changes that occur with profound interruption of breathing. The compartmentalization of the preBötC into segregated subgroups of neurons based on their connections impacts our comprehension of mechanisms that coordinate and integrate breathing with different motor and physiological behaviours. This is of fundamental importance, given that abnormal respiratory modulation of autonomic activity and orofacial behaviours have been associated with the development and progression of diseases (El-Omar et al., 2001; Huff et al., 2022; Menuet et al., 2017; Simms et al., 2009). For example, exaggerated respiratory-sympathetic modulation seems to be an important mechanism for the development and progression of hypertension (Barnett et al., 2020; Menuet et al., 2017; Moraes et al., 2014; Simms et al., 2009; Tatasciore et al., 2007). Reduced inspiratory modulation of HR is associated with the severity of diverse pathological conditions such as hypertension, heart failure, depression and anxiety (El-Omar et al., 2001; Masi et al., 2007; Thayer et al., 2012). In addition, loss of the synchronization between breathing and orofacial and oropharyngeal behaviours, such as, swallowing, has significant clinical implications, as it increases the risk of aspiration pneumonia associated with dysphagia, which can lead to death (Beard et al., 1996; Heemskerk and Roos, 2012; Huff et al., 2022).

### The impact of inhibition of preBötC neurons on orofacial motor activity is state-dependent

Nasofacial and orofacial activity are sensitive to anesthesia and animal state (Deschênes et al., 2016, 2015). Naris dilation and whisker motion are silent when animals are deeply anesthetized with ketamine/medetomidine but resume in synchrony with the inspiratory phase of the respiratory cycle, as rats recover from anesthesia (Deschênes et al., 2015). We also observed that mystacial pad activity is minimal and non-rhythmic under surgical anesthesia with ketamine/medetomidine, but resumes, with inspiratory modulation, as the animal recovers consciousness. In contrast, strong inspiratory-related mystacial pad activity occurs when surgical anesthesia is provided by urethane. In both cases, surgical anesthesia was defined by loss of the pedal withdrawal and corneal reflexes. The mechanisms by which deep ketamine and urethane anesthesia differentially affect respiratory-related motor activity are unknown. Under light ketamine anesthesia, GABAergic cells of vIRt displayed phase-locked firing patterns for protraction or retraction phases of whisking, whilst only retraction unit activity was recorded under light urethane anesthesia, suggesting that only inhibitory vIRt cells, whose activity is correlated with vibrissa retraction, are active in urethane-anesthetized rats (Deschênes et al., 2016).

Likewise, the effect of photoinhibition of preBötC→7n neurons on mystacial pad activity is both state- and anesthetic-dependent. Under urethane anesthesia, selective photoinhibition of preBötC→7n neurons silenced mystacial pad activity, whilst photoinhibition during different phases of recovery from ketamine/medetomidine anesthesia interrupted the inspiratory-related and increased the overall tonic mystacial pad activity. The reason for this substantial state-dependent effect of the preBötC→7n on mystacial pad activity remains unclear, but it has been reported in other systems. Injection of L-glutamate in the NTS induces a substantial pressor response in awake normotensive rats but a depressor response in the same rats anesthetized with urethane (Machado and Bonagamba, 1992). Photoinhibition of the C1-RVLM neurons in awake rats produces a very small depressor response (Wenker et al., 2017) when compared to that observed in anesthetized animals (Marina et al., 2011). Likewise, electrical stimulation of the central amygdala (CeA) induces a significant pressor response in conscious rats but a depressor response under anesthesia (Chiou et al., 2009). Changes in the activity of several fast ionotropic transmitter systems, including glycine, GABA, acetylcholine and glutamate, have been reported in urethane anesthesia (Accorsi-Mendonça et al., 2007; Hara and Harris, 2002). Whilst speculative, we hypothesize that under urethane, the glutamatergic preBötC→7n neurons play a major role in regulating the respiration-related mystacial pad activity, whilst the GABAergic neurons are more active in the conscious state.

Under urethane-anesthesia and following recovery from ketamine/medetomidine, inspiratory-related mystacial pad activity is clearly evident. The electrodes we used to record mystacial pad activity were inserted near the rostral tip of the snout, which is known to be the site of origin of the *nasolabialis profundus*. However, due to the large number of distinct muscles that compose the mystacial pad (Haidarliu et al., 2010), the difficulty of dissecting the rodent snout (Haidarliu et al., 2012), the substantial size of the electrode’s suture pads and the fact that the *nasolabialis profundus* electromyogenic activity was not recorded prior to the electrode implantation, it is possible that the mystacial pad activity we recorded reflects the activity of multiple muscles. Nonetheless, given that *nasolabialis profundus* is active during basal respiration, while extrinsic muscles are silent, and preferentially contracts during the sniffing and whisking state (Britto et al., 2020; Deschênes et al., 2015), we conclude that the inspiratory component of the mystacial pad activity predominantly reflects the *nasolabialis profundus* muscle contraction.

Non-selective inhibition of preBötC induced apnea and immediate interruption of the inspiratory-related mystacial pad activity, which increased in amplitude and became tonic. The inspiratory-related mystacial pad activity recovered when breathing resumed. This coordinated activity between breathing and orofacial/nasofacial activity has been described before (Deschênes et al., 2015). It was demonstrated that both naris dilation and vibrissae retraction are abolished during apnea induced by the application of ammonia to the snout, and their activity synchronously recovers with the resumption of breathing. Nonetheless, the neural mechanisms that underly these responses are still unclear. Non-selective photoinhibition of preBötC neurons may lead to inhibition of preBötC→7n, and at the same time, to inhibition of preBötC inhibitory neurons that project to parvalbumin-expressing inhibitory neurons of the vIRt neurons (vIRtPV). The vIRtPV was recently identified as the whisking oscillator (Takatoh et al., 2022), and direct projections from preBötC have been shown to reset the vIRtPV activity (Deschênes et al., 2016; Golomb et al., 2022; Takatoh et al., 2022). Moreover, since the p*ost-mortem* analysis showed that the non-selective approach transduced some vIRtPV neurons, it is also likely that the vIRtPV activity could have been directly affected by the laser delivery producing desynchronization of facial motoneurons activity and suppression of the rhythmic whisking (Takatoh et al., 2022).

### Photoinhibition of preBötC neurons modulated expiratory abdominal muscle activity

Beyond affecting inspiratory and MP activity, non-selective inhibition of preBötC also increased and produced tonic activity of the abdominal muscle that lasted for the apnea duration. The active expiratory activity of the abdominal muscles is strictly controlled by quiescent and synaptically inhibited late-expiratory neurons located in the lateral parafacial nucleus (Magalhães et al., 2021; Pagliardini et al., 2011). Our results support the evidence that preBötC is a potential source of inhibitory input to lateral parafacial. This interaction could be essential for the generation of active expiration in situations with increased respiratory demand, such as hypercapnia/acidosis (del Negro et al., 2018). Interestingly, in addition to regulating expiratory activity, it has been shown that the lateral parafacial nucleus may also play a role in coordinating nasofacial and orofacial behaviour during high chemical drive via direct projections to 7n (Britto et al., 2020). Thus, indirect projections from preBötC to 7n via lateral parafacial nucleus may also be involved with the responses on the mystacial pad activity induced by the non-selective inhibition of preBötC.

### Facial projecting preBötC neurons have functionally-relevant collateral projections

Whilst there was a clear difference in the effect of selective preBötC→7n photoinhibition on breathing, compared to non-selective preBötC photoinhibition, selective photoinhibition did have small effects on breathing, particularly under ketamine/medetomidine anesthesia and its recovery phases. With the selective approach, we observed a much smaller population of transduced neurons with a more restricted anatomical location. We did not see any transgene expression in the parvalbumin-expressing neurons of rVRG (Alheid et al., 2002; Wu et al., 2017). A similar proportion of GABAergic and glutamatergic neurons expressing GtACR2-MuGFP were found with both selective and non-selective approaches. We observed that the transduced axons of preBötC→7n neurons projected widely into multiple brainstem nuclei, including RVLM, nA, NTS, BötC, rVRG and vIRt. Collateral projections from excitatory and inhibitory preBötC neurons to premotor and motoneurons have been described before and appear to be essential for the coordination of inspiratory and expiratory activity (Koizumi et al., 2013). It was suggested that divergent axonal projections from the commissural excitatory preBötC into the HN and rVRG may be essential for initiating the inspiratory activity. On the other hand, feedforward inhibition via collateral projections from inhibitory preBötC into these same regions may contribute to the dynamic shaping of the respiratory pattern (Koizumi et al., 2013). Whilst the functional role of many of these collateral projections has not been established yet, we hypothesize that the small effect of selective photoinhibition reflects their ongoing, functionally-relevant activity. It is possible that the strict synchronization between premotor and motor neurons is not confined to respiratory motor activity, but is crucial for the coordination of respiratory, cardiovascular, and orofacial/nasofacial activity necessary for the execution of complex behaviours, such as exercise, response to stress or pain.

### Ideas and Speculation

The preBötC, initially defined as the kernel for generating inspiratory rhythm, appears to act as a master oscillator regulating other physiological functions, such as orofacial behaviours and autonomic nervous activity. Our study suggests that groups of these neurons play principal roles in these specific functions. However, the widespread collateralisation we observed along with effects of photoinhibition of the preBötC-7n on disparate motor activities raises the possibility that preBötC neurons may coordinate multiple outputs. Whether this is a function of most neurons projecting to more than one target, or a small number of neurons projecting to many targets, remains to be determined.

As has been described previously using transgenic mice (Yang and Feldman, 2018), we also observed that different subpopulations of preBötC-7n express either markers of an excitatory phenotype or an inhibitory phenotype. The idea that one output nucleus provides both excitatory and inhibitory projection to a target is intriguing. Our methods could not determine whether these different neurochemical groups might project to the same target neuron, or whether the pathways are parallel and separate. It is possible that the different groups are active under different states and enable altered coupling to the respiratory cycle. For example, under urethane anesthesia, when mystacial pad EMG is active, photoinhibition of preBötC→7n resulted in a decrease in activity. In contrast, in conscious rats, the same photoinhibition of preBötC→7n increased activity.

Exaggerated inspiratory modulation of sympathetic activity is associated with the development of hypertension(Menuet et al., 2017; Simms et al., 2009), which can increase BP variability. Interestingly, independent of the BP level, BP variability seems to be an important contributor to organ damage, such as renal dysfunction and left ventricular hypertrophy, cardiovascular disease, and poor clinical outcomes (Messerli et al., 2019). The central network underlying this autonomic dysfunction remains to be elucidated. Our previous studies suggest that respiratory input from preBotC to the catecholaminergic C1 neurons of RVLM may be involved with the increased inspiratory-sympathetic modulation in hypertensive rats (Menuet et al., 2017). In this way, methodologies that only affect inputs from preBotC to C1 neurons, such as the approach used in the present study, could shine the light on the mechanisms involved in the development of hypertension and possibly contribute to the development of targeted therapeutics used to prevent hypertension development.

## CONCLUSION

We tested the hypothesis that the preBötC might consist of separate subpopulations of neurons that project to specific nuclei to coordinate respiratory rhythmicity with different physiological behaviours, such as nasofacial activity. We showed that even when selecting just neurons projecting to a specific target, both excitatory and inhibitory neurons were transduced. Selective photoinhibition of these neurons enabled observation of the effect on nasofacial motor activity in the absence of substantial changes in respiratory, or other autonomic, activities. However, small effects on these other functions, such as diaphragm muscle activity, remain. We observed collateral axonal projections of preBötC-7n neurons to several brainstem nuclei, including the rVRG, and conclude that these are functional, active projections. This unmasks the possibility that these neurons may play a role in the complex synchronization between respiratory, cardiovascular, orofacial, and potentially other, physiological functions.

## MATERIAL AND METHODS

### Animal experiments

Experiments were conducted in accordance with the National Health and Medical Research Council of Australia’s “Guidelines to promote the well-being of animals used for scientific purposes: The assessment and alleviation of pain and distress in research animals (2008)” and “Australian code for the care and use of animals for scientific purposes” and were approved by the University of Melbourne Animal Research Ethics and Biosafety Committees (ethics ID#21396). All experiments were performed on male Sprague-Dawley (SD) rats, initially weighing 60-80g. The rats were housed in standard cages in groups of up to 4, had *ad libitum* access to standard chow and tap water, and were maintained under a 12:12h light-dark cycle in a 21⁰C temperature-regulated room at the University of Melbourne Biological Resources Facility (BRF).

### Plasmid design and generation of AAV-DIO-GtACR2-MuGFP

C-terminal fusion of a 65-amino acid trafficking motif from the voltage-gated potassium channel Kv2.1 (Lim et al., 2000) has been shown to enrich the expression of the *Gt*ACR2 in the somatic membrane of mouse cortical neurons (Mahn et al., 2018; Messier et al., 2018). We employed an analogous design in the present study to develop a *Gt*ACR2^Kv2.1^ fusion construct, with a monomeric, ultra-stable green fluorescent protein (MuGFP) tag (Scott et al., 2018) linked with four alanine residues to the *Gt*ACR2 C-terminus. To achieve Cre-dependent expression, *Gt*ACR2^Kv2.1^-MuGFP was incorporated into the pAAV-hSyn-DIO-(hCAR)off-(ChETA-mRuby2)on-W3SL plasmid (Addgene plasmid #111391; (Li et al., 2018); a gift from Prof Adam Kepecs (Cold Spring Harbor Laboratory, NY, USA)), using NheI and PacI restriction sites to replace the ChETA-mRuby2 coding sequence. The hSyn promoter was subsequently excised, and a CAG promoter sequence was ligated in its place via XbaI restriction sites. Following Cre-mediated recombination, the orientation of *Gt*ACR2^Kv2.1^-MuGFP coding sequence is reversed relative to the promoter, enabling the expression of *Gt*ACR2^Kv2.1^-MuGFP.

Co-transfection of pAAV-CAG-DIO-(hCAR)off-(*Gt*ACR2^Kv2.1^-MuGFP)on-W3SL with pDPI and pDPII plasmids (Grimm et al., 2003) into AAV293 cells (Agilent Technologies, CA, USA) preceded harvesting and iodixanol gradient purification (as described by (Zolotukhin et al., 1999) and (Ganella et al., 2013)) of AAV1/2-CAG-DIO-(hCAR)off-(*Gt*ACR2^Kv2.1^-MuGFP)on-W3SL vector – hereafter referred to as AAV-DIO-GtACR2-MuGFP. Titration of purified AAV vector was performed using quantitative polymerase chain reaction (qPCR) as described by (Ma et al., 2017) using forward (5’-CATTCTCGGACACAAACTGGAGTACAAC) and reverse (5’-GTCTGCTAGTTGAACGGAACCATCTTC) primers targeting the MuGFP coding sequence, rather than WPRE (which is replaced here by W3SL). Primers were synthesized by Bioneer Pacific (VIC, Australia). GtACR2-MuGFP, Kv2.1, and CAG sequences were synthesized by GenScript (NJ, USA). Restriction enzymes and T4 DNA ligase were sourced from New England Biolabs (VIC, Australia) and Promega (NSW, Australia), respectively, and used according to the manufacturer’s recommendations.

### Other viruses

Non-selective expression of GtACR2^Kv2.1^-MuGFP in the preBötC was achieved by the injection of a mixture containing AAV-DIO-GtACR2-MuGFP (2.04x10^11^GC/ml) and AAVDJ8-CBA-Cre-TdTomato-WPRE (1.14x10^12^VP/ml) into preBötC. Selective expression of GtACR2-MuGFP-Kv2.1 in preBötC neurons that project to the 7n was obtained by injections of AAV-DIO-GtACR2-MuGFP into the preBötC and the retrograde pseudotyped AAVrg-EF1α-mCherry-IRES-Cre (titre 8x10^12^ GC/ml; Addgene # 55632-AVVrg) into the 7n.

### Retrograde Labelling Cholera toxin subunit B (CTB)

Retrograde labelling of preBötC→7n neurons was achieved by either the injection of AAVrg-CAG-GFP (1.29x10^13^VP/ml) or Alexa Fluor 647-conjugated cholera toxin subunit B (CTB; Sigma, C34778) into 7n. Retrograde labelling of preBötC→RVLM neurons was achieved by injection of either AAVrg-EF1α-mCherry-IRES-cre or Alexa Fluor 555-conjugated CTB (Sigma, C22841) into RVLM. CTB was dissolved in PBS and injected at a concentration of 0.5%.

### Microinjection into the brainstem

Animals were anesthetized with an intraperitoneal injection of ketamine (75 mg/kg, i.p., Lyppard, Dingley, Australia) and medetomidine (0.5 mg/kg, i.p., Pfizer Animal Health, West Ryde, Australia). Eye moisture was maintained by the application of a hydrating gel (POLY VISC® Eye Ointment, Alcon). The surgical field was shaved and disinfected with 80% ethanol and chlorhexidine. Surgery was initiated once a deep surgical level of anesthesia was obtained, as evidenced by loss of the pedal withdrawal and corneal reflexes. Throughout the protocol, body temperature was maintained at 37.5⁰C with a heating pad (TC-1000 Temperature controller, CWE Inc.) that was covered with an autoclaved non-absorbent pad. Rats were then placed in a stereotaxic frame with the nose ventro-flexed (incisor bar -15 mm; RWD Life Science). Extracellular recordings of multiunit activity were used to functionally map the preBötC, as previously described (Menuet et al., 2020). For all injections, the pipette was 20° angled in the caudal-rostral axis, the tip pointing forward. For injections into preBötC, the pipette was descended into the brainstem 1.5 mm lateral to the midline and the injections were made at the most rostral point at which vigorous inspiratory-locked activity was isolated (typically 0.2 ± 0.1 mm rostral, 1.6 ± 0.4 mm ventral to the *calamus scriptorius*). The injections into preBötC were made bilaterally, and a picospritzer (World Precision Instruments, Sarasota, USA) was used to microinject the virus (50 nl over 5 min on each side).

Antidromic field potentials, elicited by stimulating the mandibular branch of the facial nerve, were used to map the dorsal, caudal, ventral, and lateral edges of the facial nucleus, allowing precise targeting of the RVLM and the facial nucleus, as described before (Menuet et al., 2017). Briefly, an electrode was positioned so that this gently touched the mandibular branch of the facial nerve. An electrical current (2.0 ± 0.5 mA) was applied in order to stimulate the nerve. Click or tap here to enter text.A picospritzer (World Precision Instruments, Sarasota, USA) was used to microinject CTB or AAVrg (40 nl per injection site over 5 min) into the RVLM immediately caudal to the facial nucleus. Injections were made bilaterally, 1.6 mm lateral to the midline at the caudal edge of the facial nucleus and 3.5 ± 0.4 mm ventral to the *calamus scriptorius*. These injections were made in 2 rostrocaudal levels separated by 300 µm; the most rostral being at the caudal edge of the facial nucleus. A picospritzer was also used to microinject AAVrg or CTB (40 nl per injection site over 5 min) into the facial nucleus. The injections were made bilaterally and targeted the lateral and dorsal-lateral borders of the facial nucleus (1.9 lateral to the midline, +1.4 and +1.1 rostral and -2.9 ± 0.2 mm ventral to the *calamus scriptorius*). At the end of the surgery, the neck muscles and the cheek skin were sutured with sterile absorbable polyglycolic acid 4/0 (Surgicryl) sutures using simple interrupted stitches.. Subcutaneous injection of non-steroidal analgesic (meloxicam 1 mg/kg, s.c., Metacam, Lyppard) and 1 ml of warmed Hartmann’s solution for fluid replacement was performed just before the intraperitoneal administration of atipamezole (1 mg/kg, i.m., Antisedan, Pfizer Animal Health, West Ryde, Australia) to reverse the effect of anesthesia. Postoperative analgesia with meloxicam and buprenorphine (0.025 mg/kg, s.c., Schering-Plough, USA) was maintained during the next 48 hours, and rats were monitored for any signs of surgical complications and weighed every day for 14 days.

### Optogenetic experiments in urethane-anesthetized rats

Following the 3-4 weeks of recovery from stereotaxic surgery, anaesthesia was induced by inhalation of isoflurane (Rhodia Australia Pty. Ltd.) in an enclosed chamber. Once anesthetized, rats were transferred to the surgical bench, where anaesthesia was maintained with 2-2.5 % isoflurane, delivered in oxygen using a SomnoSuite low-flow anesthesia delivery system (Kent Scientific). Body temperature was maintained at 37.5 °C with a TC-1000 heat pad (CWE Inc.). A pulse oximeter probe was placed on a paw, and the body temperature was monitored using a rectal probe attached to the SomnoSuite anesthesia equipment. For diaphragm electromyography (dEMG) recordings, a lateral abdominal incision was made, and two nylon-insulated stainless-steel wire electrodes (0.25 mm insulated diameter) ending with suture pads (0.7 × 1.0 × 3.2 mm, Plastics One, VA, USA) placed in the costal diaphragm, 3–4 mm apart. For abdominal (Abd) EMG recordings, a lateral transverse incision was made in the abdomen region, and two silver wire electrodes (0.635 mm insulated diameter, A-M Systems) were implanted in the oblique abdominal muscles, 4-5 mm apart. For mystacial pad EMG, an incision was made along the midline of the nose, and two silver wires (0.38 mm insulated diameter, A-M Systems) were implanted into the mystacial pad near the rostral area of the snout – approximately between row C and D and columns 5-7, 3-4 mm apart. The femoral artery and vein of the left leg were cannulated for measurement of arterial pressure and drug administration, respectively (PE10 tubing (ID 0.28 x OD 0.61 mm) connected to PE50 (0.17 x 1.45 mm)). Isoflurane anaesthesia was gradually replaced by 1.2 g/kg intravenous urethane, following which rats were tracheotomized, and oxygen (100%) was directed over the tracheotomy cannula. The arterial catheter was connected to a Statham Gould (P23 Db) pressure transducer, and the dEMG, AbdEMG and mystacial pad EMG electrode wires were coupled to an amplifier (10 Hz- 5 kHz band-pass filter, 5 kHz sampling rate, Model 1700 Differential AC amplifier, A-M systems, Sequim, WA, USA). Arterial pressure and electromyography were recorded using Spike2 software (Cambridge Electronic Design). Rats were transferred to a stereotaxic frame (incisor bar +3 mm; RWD Life Science). To perform the optogenetic experiments, the optical fibres (200 μm diameter, 0.22 NA, Ø1.25 mm, RWD Life Science) were secured to a manipulator on the stereotaxic frame and lowered through the dorsal surface of the skull through burr holes made bilaterally in the occipital bone centred approximately 1 mm rostral to the occipital suture, 1.8 ± 0.2 mm lateral from the midline. Output intensity from the optical fibres was determined using a PM100D Meter (Thorlabs, NJ, USA). The strongest physiological responses to *Gt*ACR2 stimulation (50 Hz, 5 ms pulse, 15-20 mW) were found at 7.5 ± 0.5 mm ventral to the surface of the skull.

### Optogenetic experiments in conscious rats

Following the 3-4 weeks of recovery from the virus injection, animals were anesthetized with an intraperitoneal injection of ketamine (75 mg/kg) and medetomidine (0.5 mg/kg). Eye moisture was maintained by the application of a hydrating gel (POLY VISC® Eye Ointment, Alcon). The surgical field was shaved and disinfected with 80% ethanol and chlorhexidine. Surgery was initiated once a deep surgical level of anesthesia was obtained, as evidenced by loss of the pedal withdrawal and corneal reflexes. Throughout the protocol, body temperature was maintained at 37.5⁰C with a heating pad (TC-1000 Temperature controller, CWE Inc.) covered with an autoclaved non-absorbent pad. Then, the rats underwent the implantation of a 6-channel pedestal with a back-mount (Type 363 components, PlasticsOne, USA) that contains percutaneous electrodes (2 × cardiac electrocardiograms (ECG), 2 × dEMG and 2 x mystacial pad EMG) and remained externalized for tethering. The electrodes have socket-pin connections at the pedestal and stainless-steel suture pads (∼0.5 mm^2^) for electrical recording (PlasticsOne, USA). For implantation of the sterilized multi-lead electrode pedestal, a transverse incision was made on the dorsal surface, approximately 1 cm below the rat’s shoulder blades, and a subcutaneous tunnel opened between the two incisions. The two ECG and dEMG electrodes were placed according to previous studies (Butler et al., 2021; O’Callaghan et al., 2020). The mystacial pad EMG electrodes were fixed with a non-absorbable suture into the mystacial pad near the rostral area of the snout – approximately between row C and D and columns 5-7, 3-4 mm apart. The skin incision was closed with sterile absorbable polyglycolic acid 4/0 (Surgicryl) sutures and the multi-lead electrode pedestal was sutured with sterile non-absorbable surgical sutures (4-0 Supramid)) into place between the scapulae. All electrodes were tunnelled to the pedestal and connected to it. At this point, the multi-lead electrode pedestal was connected to an amplifier (10 Hz-5 KHz band-pass filter, 5 kHz sampling rate, Model 1700 Differential AC amplifier, A-M systems, Sequim, WA, USA). The signal was recorded and integrated using Spike2 version 9.0 software (Cambridge Electronic Design).

Following the implantation of the electrodes, rats were transferred to a stereotaxic frame (incisor bar +3 mm; RWD Life Science). To perform the optogenetic experiments, the optical fibres (200 μm diameter, 0.22 NA, Ø1.25 mm, RWD Life Science) were secured to a manipulator on the stereotaxic frame and lowered through the dorsal surface of the skull through burr bilateral holes made in the occipital bone centred approximately 1 mm rostral to the occipital suture, 1.8 ± 0.2 mm lateral from the midline. Output intensity from the optical fibres was determined using a PM100D Meter (Thorlabs, NJ, USA). Physiological responses to GtACR2 stimulation were investigated at 0.5 mm increments moving ventrally from the surface of the skull. The strongest physiological responses to *Gt*ACR2 stimulation (50 Hz, 5 ms pulse, 15-20 mW) were typically observed at 7.5 ± 0.5 mm ventral, and it was where the optical fibers were fixed with dental cement (Pattern Resin LS, GC America INC, IL).

The rat was transferred to a recording cage, and the effects of preBötC photoinhibition on dEMG, ECG and mystacial pad EMG were evaluated under three different states: when the animals were still under ketamine/medetomidine anesthesia, during the very early stage of regaining consciousness, a few min after the anesthesia was reversed with atipamezole and 1-2 h after the atipamezole injection when the animal was completely awake and had recovered from anesthesia.

### Immunohistochemistry

At the end of the urethane-anesthetized and conscious experiments, animals were perfused transcardially with 0.1M phosphate-buffer saline (PBS) (1 ml/g of b.w.) followed by 4% formaldehyde (FA) in 0.1 M PBS using a peristaltic pump (Masterflex L/S Drive System, Cole-Parmer, Vernon Hills, Illinois, USA). The brains were removed, post-fixed in 4% PFA for 12 h at 4°C, and then immersed in 20% sucrose at 4°C until processing. Brainstems were frozen at -20°C, and four series of coronal sections (40 µm thickness) were cryostat sectioned. Sections were placed directly into cryoprotectant in a 24-well plate and stored at −20°C until processing. Fluorescence immunohistochemistry was performed as previously described (Bassi et al., 2022; Menuet et al., 2020; Ngo et al., 2020). Primary antibodies used were chicken anti-GFP (1:5000, Aves Labs Inc, Davis, CA, USA, Cat#: GFP-1010), mouse anti-TH (1:5000, Merck-Millipore, Bayswater, VIC, Australia, Cat#: MAB318), rabbit anti-parvalbumin (1:5000, Abcam, Melbourne, VIC, Australia, Cat#: ab11427), Goat anti-ChAT: (1:1000, Chemicon-Merck, Bayswater, Australia, Cat#: AB144P), rabbit anti-dsRed: (1:5000, Takara Bio, Clontech, Australia, Cat#: 632496), goat anti-mCherry (1:5000, Sicgen, Cantanhede, Portugal; Cat#: AB0040-200).

All secondary antibodies were purchased from Jackson ImmunoResearch Laboratories, Inc, West Grove, Pennsylvania, USA, and were used at either 1:500. The following secondary antibodies were used: Alexa488-conjugated donkey anti-chicken (Cat#: 703545155), Cy3-conjugated donkey anti-rabbit (Cat#: 711165152), Cy3-conjugated donkey anti-goat (Cat#: 705165003), Cy5-conjugated donkey anti-mouse (Cat#: 715175151), and Cy5-conjugated donkey anti-goat (Cat#: 705175003) and biotin-SP-conjugated donkey anti-goat (Cat#: 705065147). Streptavidin-Marina blue (1:200, Cat#: S11221) was purchased from Invitrogen-Thermo Fisher Scientific, Scoresby, VIC, Australia.

### In situ hybridisation - RNAscope

Brains from SD rats with the selective and non-selective expression of GtACR2-MuGFP were used to perform this analysis. Briefly, brains were freshly extracted (n = 3 per group) and frozen in isopentane and dry ice, then stored at -80°C. Brainstems were cryostat sectioned (16 µm), collected and mounted onto Superfrost-Plus slides (Thermo Scientific™). Frozen sections were fixed in 4% PFA for 16 min at 4°C, rinsed twice in DEPC-treated 1X PBS for 1 min each, then dehydrated in 50%, 70% and 100% ethanol solutions for 5 min each. Dehydrated sections were then stored in 100% ethanol overnight at -20°C. The following day, slides were air-dried at room temperature (RT), and hydrophobic barriers were created using ImmEdge hydrophobic PAP pen(Vector Laboratories, Burlingame, CA, USA). Sections were incubated with Protease IV (ACD, CA, USA; Cat #322336) for 20 min at RT, rinsed twice with DEPC-treated 1XPBS for 1 min each before incubating with the probe mixture for 2 h at 40°C. The probe mixture consisted of probes against VGAT (SLC32A1-ACD;Cat #ADV424541-C2) and VGluT2 (Slc17a6, ACD; Cat #317011-C3) and C1 diluent (ACD, Cat #ADV300041). Signal amplification was achieved in accordance with the RNAscope multiplex fluorescence v1 kit manufacturer instructions (ACD, Cat #320851).

Immediately following the RNAScope protocol, the immunofluorescence protocol for MuGFP was performed as previously described (Bassi et al., 2022) and nuclei DAPI stained. 4’,6-diamidine-2-phenylindole dihydrochloride And coverslipped using ProLong™ Gold Antifade Mountant (Invitrogen by Thermo Fisher Scientific).

### Imaging

Imaging was performed at the Biological and Optical Microscopy Platform, University of Melbourne as previously described(Bassi et al., 2022). Lower power images were taken using epifluorescence microscopy (Zeiss Axio Imager D1 microscope with a Zeiss AxioCam MR3 camera and EC Plan-NeoFuar 10x/0.3 air objective, Carl-Zeiss, North Ryde, NSW, Australia). High-resolution images were captured on Zeiss LSM880 confocal laser scanning microscope with Airyscan detectors (32 GaAsP PMT array) at 20X using a Plan-Apochromat 20x/0.8 M27 objective (Carl-Zeiss, North Ryde, NSW, Australia) and ZEN 2.3 (black edition) imaging software. Z-stack images were taken with 0.4 µm z-intervals, and at least 50% overlap between optical slices obtaining 40-50 images over 15-30 µm thickness. Tile scan was performed with 10% overlap between neighbouring tiles to image the entire distribution of labelled axonal fibers in the section. High-resolution images were also captured with high magnification using a Plan-Apochromat 63x/1.4 Oil DIC M27 objective.

### Mapping of reporter expression

Whilst GtACR2-MuGFP expression was examined across all animals, the maps of the distribution were generated from 7 and 5 representative animals of non-selective and selective expression of GtACR2-MuGFP, respectively. To build the maps of distribution, the images containing GtACR2-MuGFP expression were selected and transformed in binary images according to a previously determined protocol (Bassi et al., 2022). The traced bitmap image was overlayed on the corresponding coronal preBötC schematic diagram (Paxinos and Watson, 2006) using the original fluorescence image as a guide. The total number of MuGFP immunofluorescent cells was determined either manually or using a macro every 160 µm. The quantification of co-localization among GtACR2-MuGFP, VGAT and VGlut2 expression was made manually each ∼200 µm. The neuron was considered as positive only if it contained at least eight dots of transcripts for the mRNA co-localized with a nucleus labeled with DAPI. The percentage of double (MuGFP/VGlut2 or MuGFP/VGAT) labelled neurons was calculated in relation to the total number of MuGFP cells that were quantified bilaterally.

### Data analysis and statistics

Analysis of the muscle activities was performed on rectified and smoothed signals (time constant of 50 ms). dEMG activity was evaluated by its burst amplitude (DIA, V) and frequency (fR, breaths/min). The DIA amplitude, fR, BP and HR values were extracted from Spike2 at the sample rate of 1 Hz and were graphically represented as absolute values. These values were taken in three different periods: during baseline (15 seconds before the beginning of photoinhibition), during the 15-30 seconds of photoinhibition and during the recovery period (15 seconds post-photoinhibition). SpikeSorting analysis (Spike2 software, Cambridge Electronic Design) was also employed to evaluate to compare the temporal activity of MP during baseline and during the photoinhibition.

The function “modulus’ of Spike2 was used to extract the areas under the curve of the raw signal of dEMG, AbdEMG and mystacial pad EMG activities. The AUC values and the changes in SAP, HR and fR were extracted from three different periods: during baseline (5 respiratory cycles before the beginning of the photoinhibition), during the photoinhibition and during the recovery (5 respiratory cycles after the end of the photoinhibition). The photoinhibition period was further divided into two periods: P1 (extracted during the 5 first respiratory cycles from the beginning of photoinhibition or, in the case where apnea was present, was collected the same extension as the 5 respiratory cycles during baseline) and P2 (5 respiratory cycles after the breathing resumption).

All sets of data were submitted to the normality test (SigmaPlot v11, CA, USA). In cases where the set of data failed to pass the normality test, the data was assessed by the nonparametric Mann-Whitney test with multiple comparisons using the Bonferroni-Dunn method (GraphPad Prism9, La Jolla, USA). If the data passed the normality test, two-way ANOVA followed by Bonferroni *post hoc*. Statistical analysis of the period P1 was performed using either the unpaired t-test, if the set of the data passed the normality test or the non-parametric Mann-Whitney U test.

All data are described as the mean [95% confidence interval (CI)] and were graphically represented as mean ± 95% CI using GraphPad (GraphPad Prism9, La Jolla, USA). Differences were considered significant at p<0.05.

## Acknowledgements

The authors acknowledge the facilities and technical assistance of the Biological Optical Microscopy Platform (University of Melbourne) for the confocal microscopy images.

## FIGURE OF LEGENDS

**Figure 1 – figure supplement 1:**
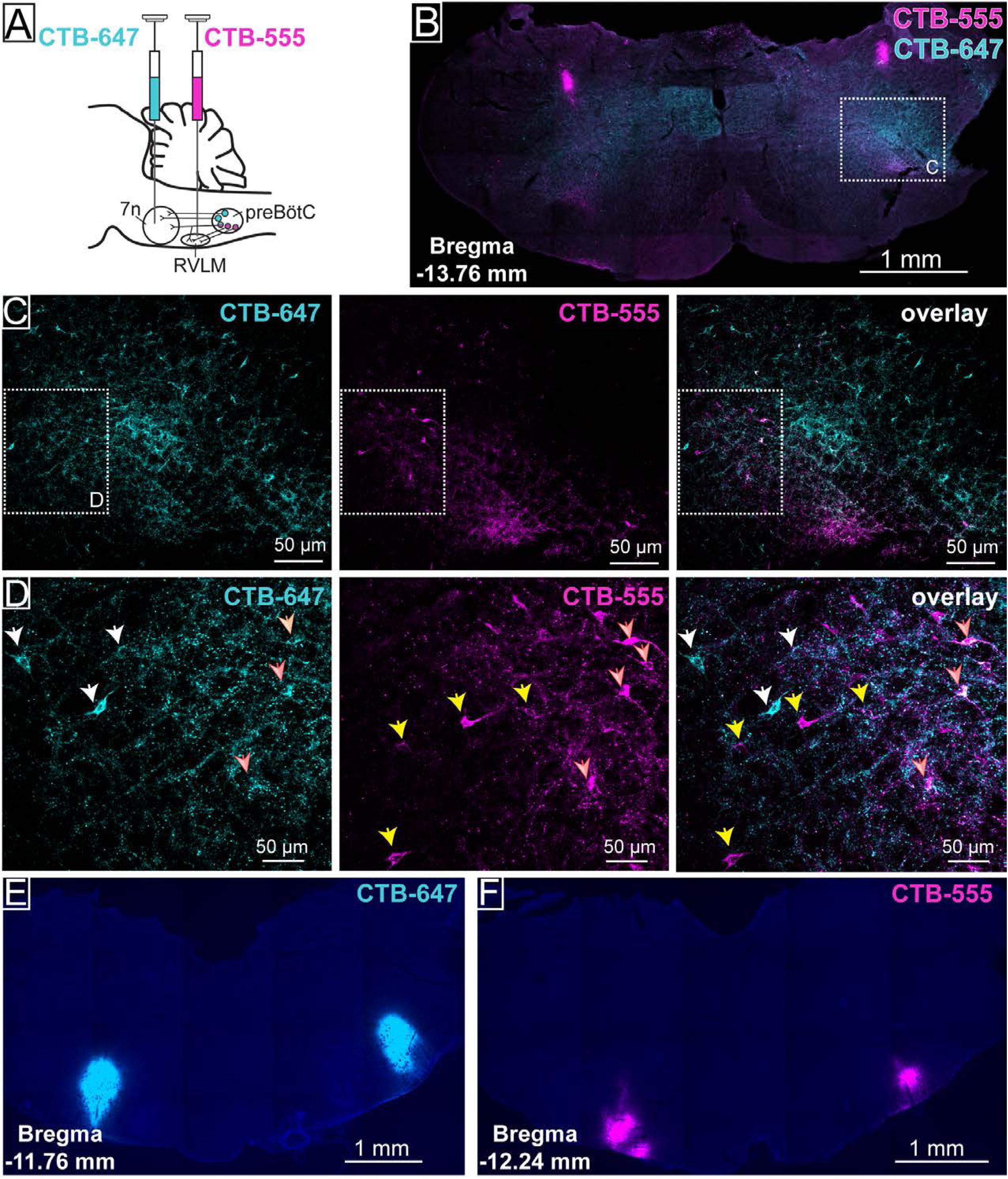
Distinct preBötC neurons project to facial nucleus (7n) and rostral ventrolateral medulla (RVLM). (A) Schematic diagram showing the dual retrograde anatomical tracer protocol. (B) Coronal section showing preBötC neurons labelled with cholera toxin B (CTB)-555 and/or CTB-647. (C) Higher magnification confocal image of preBötC neurons in the hashed box in (B). (D) Higher magnification of the inset in (C) showing preBötC neurons that project only to 7n (white arrows), only RVLM (yellow arrows), or to both 7n and RVLM (orange arrow with white). Images showing the injection site in 7n (E) or RVLM (F).

**Figure 2 – figure supplement 1:**
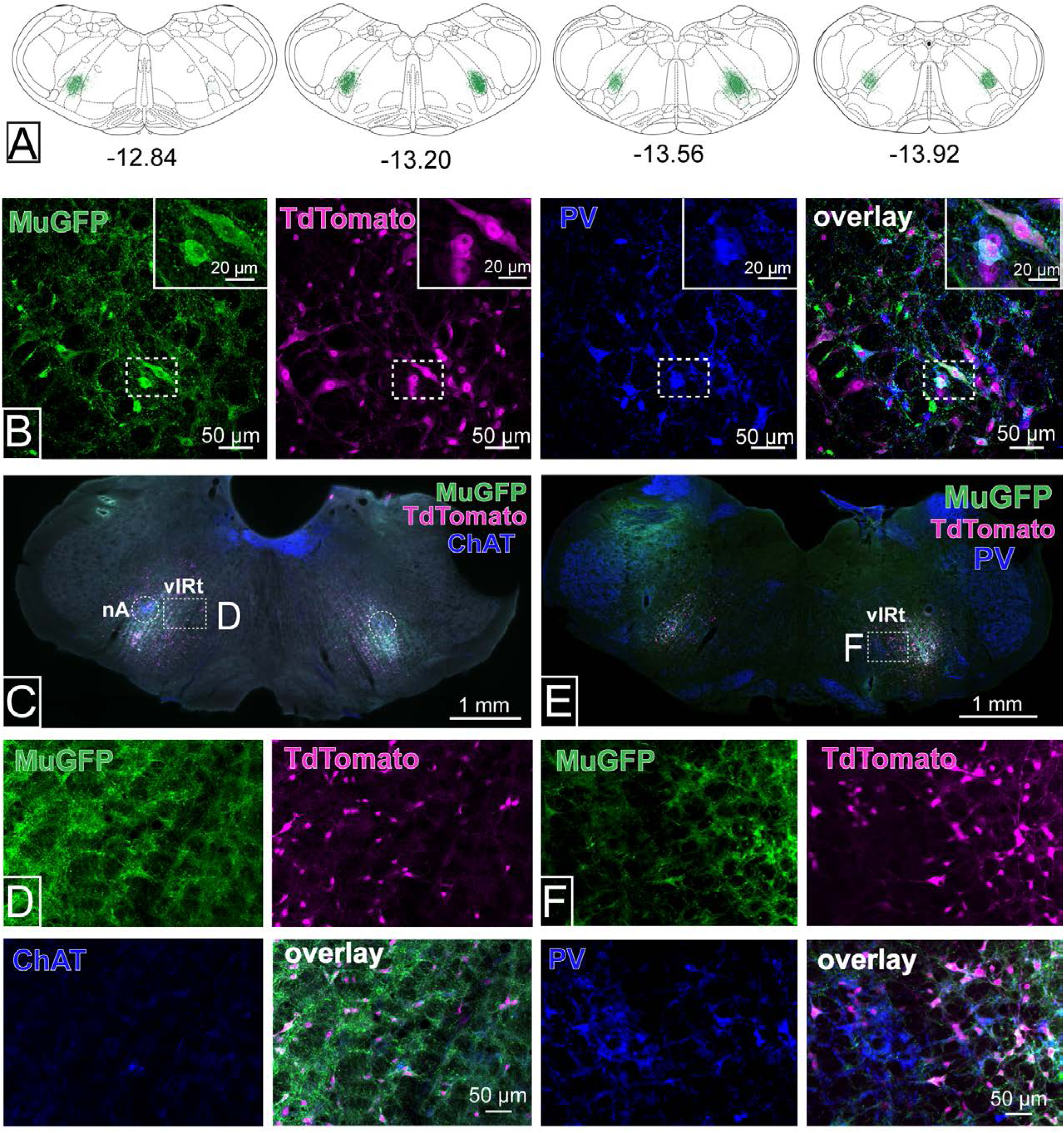
Expression of GtACR2-MuGFP in rats co-injected with AAV-DIO-GtACR2-MuGFP and AAVDJ8-Cre-TdTomato in the preBötC. (A) Schematic coronal sections of the rat medulla, based on the atlas of Paxinos and Watson, 7ed, showing a heat map to depict the distribution of GtACR2-MuGFP along the rostral to the caudal extension of the preBötC with levels relative to Bregma (mm) from a cohort of urethane-anesthetized rats (n = 7). The distribution in each animal is shown, with increased intensity of the green depicting overlapping areas of transduction. Sections rostral or caudal to these levels had no labeling in any animal; (B) Viral transgene expression in parvalbumin (PV) expressing neurons of the rostral ventral respiratory group (rVRG). (C, E) Viral transgene expression in the vibrissa intermediate reticular nucleus (vIRt), which is located medially to nucleus ambiguus (nA) and depicted by a hashed boxes, in combination with immunohistochemistry for choline acetyltransferase (ChAT) (C,D) and parvalbumin (PV) (E,F). The hashed circles in (C) show the nA.

**Figure 2 – figure supplement 2:**
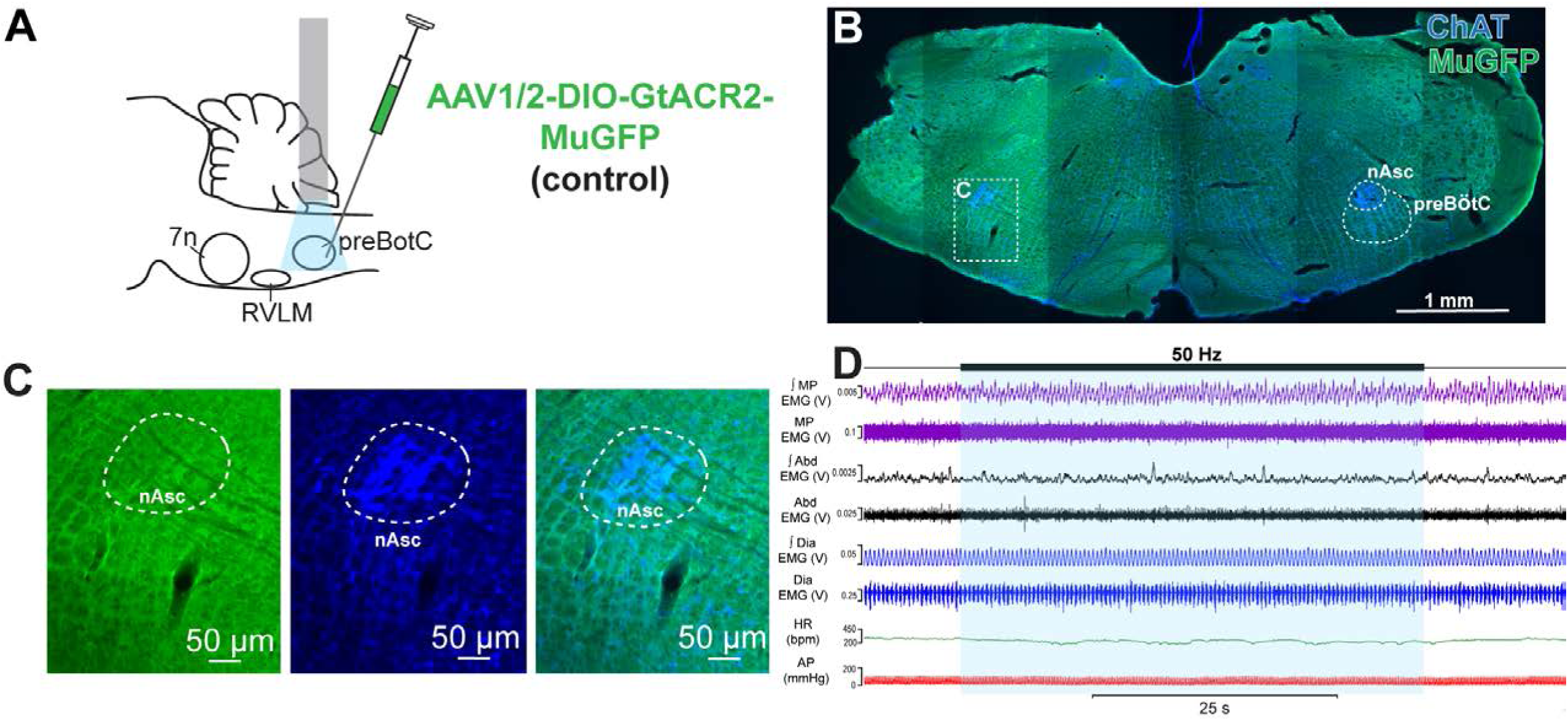
Control experiments with injection of AAV-DIO-GtACR2-MuGFP alone into preBötC. (A) Schematic diagram showing the injection protocol. (B) Coronal section showing that injection of AAV-DIO-GtACR2-MuGFP did not induce transgene expression in the preBötC. Immunoreactivity for choline acetyltransferase (ChAT) is used to show the position of the preBötC. (C) Higher magnification confocal image showing the absence GtACR2-MuGFP expression in the preBötC. (D) Representative trace showing the integrated (∫) and raw mystacial pad (MP) EMG, ∫ and raw abdominal muscle (Abd) EMG, ∫ and raw diaphragm (Dia) EMG, heart rate (HR), and arterial pressure (AP). The period of bilateral laser delivery into the preBötC is shown by the blue box. No changes in any of these parameters were observed in control rats. Abbreviation: nAsc: subcompact formation of the nucleus ambiguus.

**Figure 3 – figure supplement 1:**
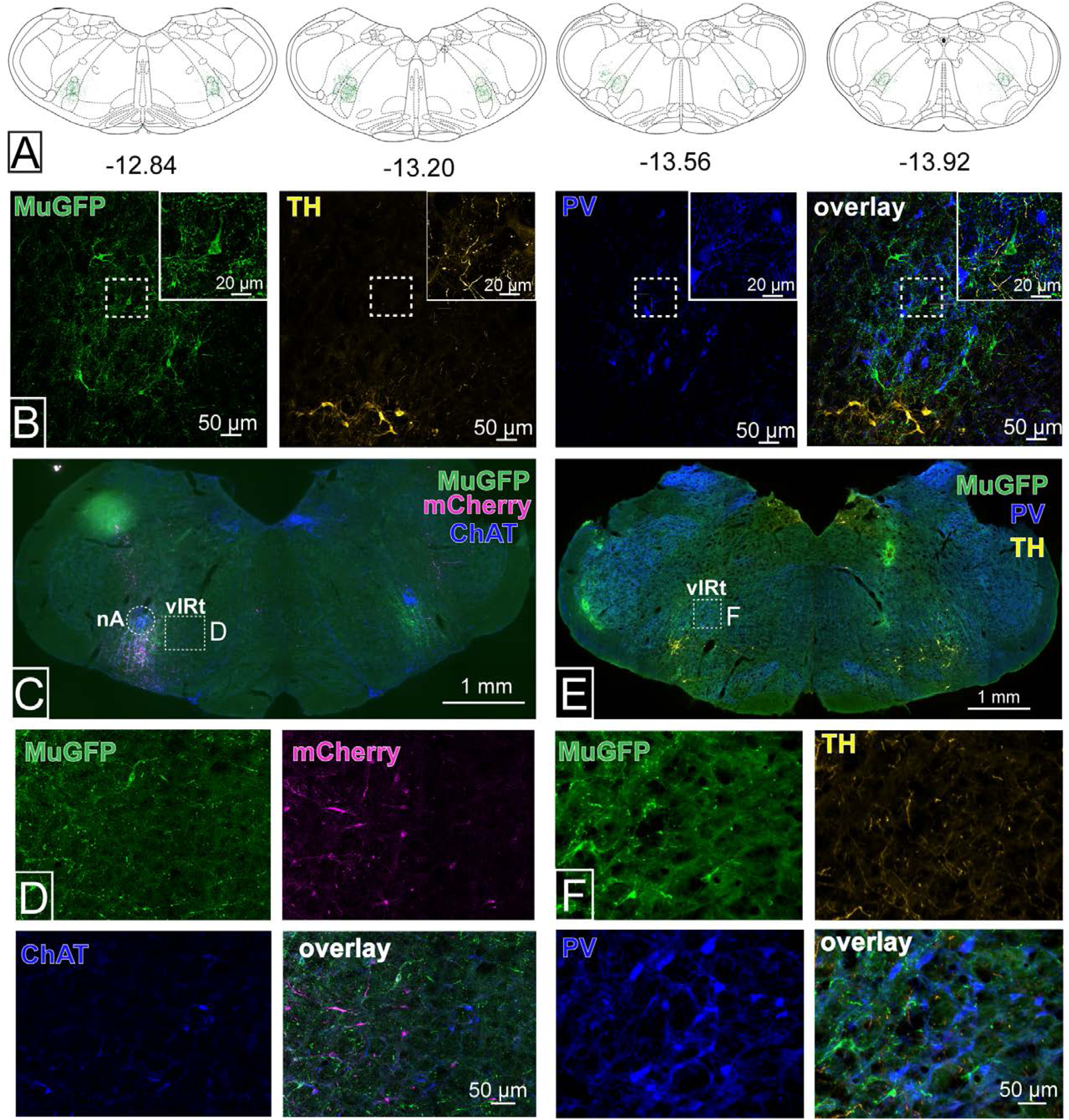
Expression of GtACR2-MuGFP in selective preBötC→7n transduced rats. (A) Schematic coronal sections of the medulla oblongata, based on the atlas of Paxinos and Watson, 7ed, showing heat maps of the distribution of GtACR2-MuGFP along the rostral-caudal extent of the preBötC with levels relative to Bregma (mm) (n = 5 from the urethane-anesthetized cohort). The distribution in each animal is shown, with increased intensity of the green depicting overlapping areas of transduction. Sections rostral or caudal to these levels had no labeling in any animal. (B) GtACR2-MuGFP expression occurs in neurons intermingled with, but not co-expressing with, parvalbumin (PV) expressing neurons of the rostral ventral respiratory group (rVRG). Immunohistochemistry for tyrosine hydroxylase (TH; yellow) shows the ventrolateral medulla. (C-D) GtACR2-MuGFP expression in the vibrissa intermediate reticular nucleus (vIRt), which is located medially to nucleus ambiguus (nA), defined by expression of choline acetyltransferase (ChAT). (E-F) Parvalbumin-immunoreactive neurons of the vIRt are intermingled with, but separate from, transduced preBötC→7n neurons, and their processes.

**Figure 7 – figure supplement 1:**
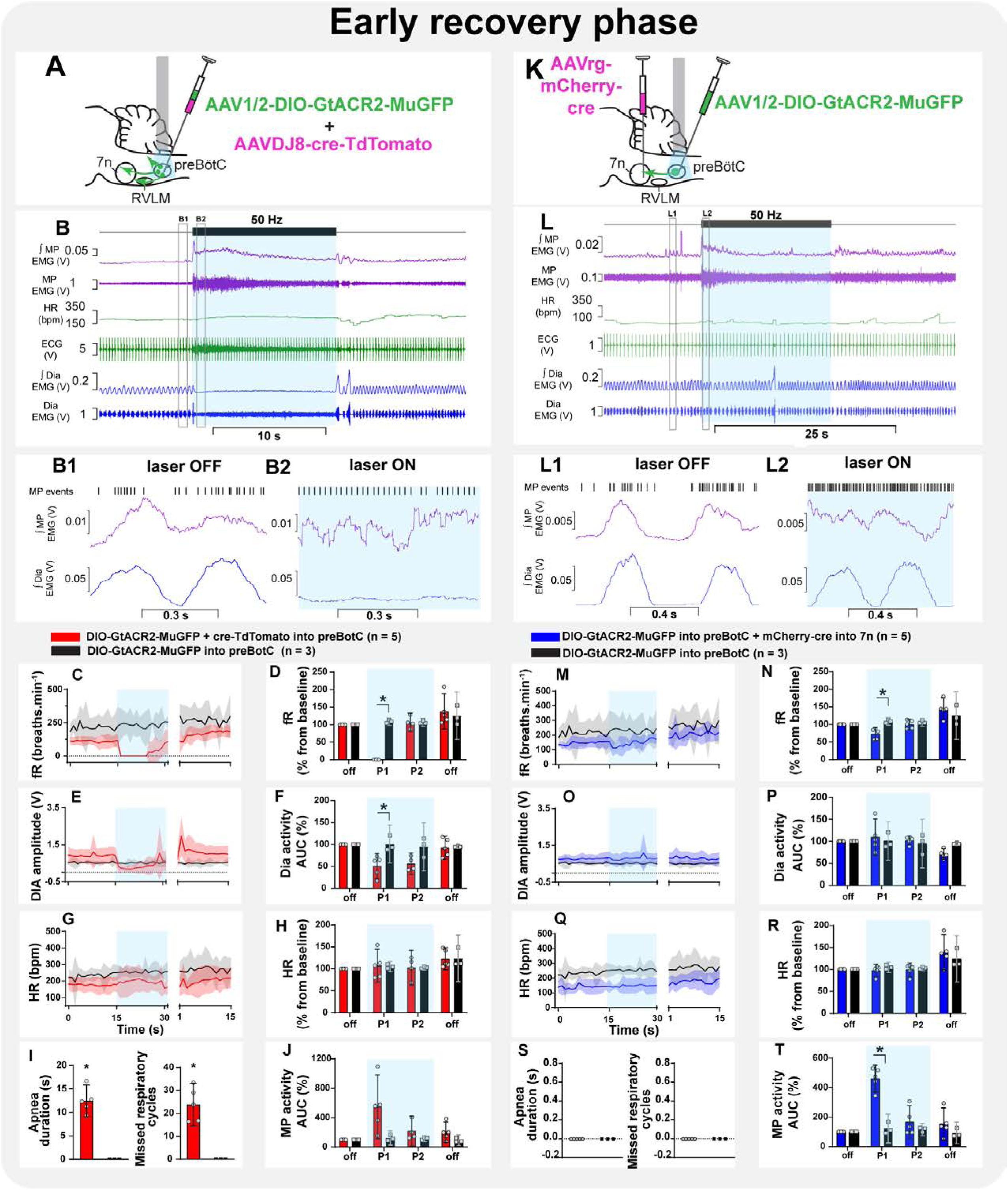
Effect on non-selective and selective inhibition of preBötC on mystacial pad activity under ketamine/medetomidine anesthesia. Schematic diagrams showing the injection protocols for non-selective transduction of preBötC neurons (A) and selective transduction of preBötC→7n neurons (K). Representative trace showing the integrated (∫) and raw mystacial pad (MP) EMG, heart rate (HR), ECG, and ∫ and raw diaphragm (Dia) EMG in ketamine/medetomidine-anesthetized rats with non-selective (B) and selective (L) transduction. The period of photoinhibition is highlighted with the blue box. Higher resolution recordings of the hashed boxes showing that MP activity was minimal and non-rhythmic during baseline (B1, L1), and not affected by the apnea in non-selective transduced rats (B2). Photoinhibition of preBötC→7n neurons had minimal effects on ∫ Dia EMG and no effect on ∫ MP EMG. Group data showing mean (solid line) and 95% confidence intervals for (C and M) respiratory frequency – fR, (E and O) Dia amplitude, (G and Q) and HR (bpm) before, during and after photoinhibition in non-selective (red), selective GtACR2 expressing (blue) and control (black) rats. Histograms showing group data for the effect of photoinhibition of preBötC on respiratory and cardiovascular parameters (D and N) fR, (F and P) Dia amplitude, (H and R) HR, (I and S) apnea duration and number of missed respiratory cycles, and (J and T) MP activity.; P1 and P2 refer to the initial period of photoinhibition. Group data are presented as mean ± 95% CI; unpaired t-test or nonparametric Mann-Whitney test with multiple comparisons using the Bonferroni-Dunn method, *p<0.05. The photoinhibition of preBötC neurons is depicted by blue shading. **Source data 1.** Source data and statistics for Figure 7 - figure supplement 1

